# The soil emergence-related transcription factor PIF3 regulates root penetration by interacting with the receptor kinase FER

**DOI:** 10.1101/2023.02.02.526371

**Authors:** Fan Xu, Jia Chen, Yingbin Li, Shilin Ouyang, Yirong Wang, Xianming Fang, Kai He, Feng Yu

## Abstract

The cotyledons of etiolated seedlings from terrestrial flowering plants must emerge from the soil surface, while roots must penetrate the soil to ensure plant survival. We show here that the soil emergence-related transcription factor PHYTOCHROME-INTERACTING FACTOR 3 (PIF3) regulates root penetration via transducing external signals perceived by the receptor kinase FERONIA (FER) in *Arabidopsis thaliana*. The loss of FER function in the *fer-4* mutant resulted in a severe defect in root penetration into hard soil or medium. Single-cell RNA-seq profiling of roots revealed a distinct cell clustering pattern, especially for root cap cells, and revealed PIF3 as one of FER-regulated transcription factor. Biochemical, imaging, and genetic experiments confirmed that PIF3 is required for root penetration into soil. Moreover, FER interacted with and stabilized PIF3, which then modulated the expression of mechanosensitive ion channel and the sloughing of outer cells in the root cap. Based on these findings, we propose a novel mechanism of soil penetration by plant roots.

## Introduction

The seeds from angiosperms often start their life cycle in subterranean darkness and must synchronously modulate both their upward emergence from soil and their downward penetration into the soil. To protect the fragile shoot apical meristem (by nonexpanded cotyledons) and root meristem (by the root cap) from mechanical injury by soil particles, seedlings employ multiple systems to maintain cellular integrity, including mechanosensitive ion channels and the cell wall integrity (CWI) pathway. Light and ethylene signaling also regulate soil emergence by recruiting specific components. For example, the central repressor of light signaling CONSTITUTIVELY PHOTOMOR PHOGENIC1 (COP1) is critical for sensing soil depth above seedlings by regulating the abundance of ETHYLENE-INSENSITIVE 3 (EIN3), a master transcription factor in the ethylene pathway^1-3^. The transcription factor PHYTOCHROME-INTERACTING FACTOR 3 (PIF3) pathway is activated in etiolated cotyledons to ensure a successful soil emergence^4^. However, while several studies have focused on the inability of the root to penetrate a substrate, it remains unknown whether soil emergence and root penetration shared common components.

Roots forage the soil for nutrients to sustain growth. As they elongate through the soil, roots can encounter belowground obstacles caused by drought and/or heterogeneous soil composition, which results in a lower growth rate, leading to reduced crop yields for the aboveground plant tissues^5^. The root tip is thought to perceive the mechanical resistance of the soil and integrate this information into a coordinated response^6,7^. The root tip consists of the root cap, the root apical meristem, and the quiescent center. The root cap, comprising the columella and lateral root cap cells^7^, offers physical protection to the root apical meristem against mechanical shearing during root penetration into the soil. In addition, root hairs provide an anchor for the root tip and assistance for emerging from empty spaces in the soil and for penetrating into bulk soil^8^. Cortical cells also contribute to root penetration^9^. Root anatomical studies have shown that smaller cells in the outer cortical region in maize (*Zea mays*) roots are associated with increased root penetration into hard soil layers by stabilizing the root against compression and reducing the risk of buckling and collapse^10^. A multiseriate cortical sclerenchyma in maize and wheat (*Triticum aestivum)* improves root tensile strength and increases penetration ability in compacted soils^11^. During soil penetration, root cells must maintain their mechanical integrity, which is determined by the CWI pathway and mechanosensitive ion channels. *Catharanthus roseus* receptor-like kinase 1-like proteins (*Cr*RLK1Ls) participate in the perception of CWI and modulate root intracellular responses. One member from the *Cr*RLK1Ls family, FERONIA(FER), is involved in CWI signaling^12,13^, as FER can be activated by the signals of pectin induced by growth and/or cell wall damage^14-17^ and small peptides Rapid Alkalinization Factor 1 (RALF1)^18^ and RALF23^19^. The FER homologs ANXUR1 (ANX1), ANX2, Buddha’s Paper Seal 1 (BUPS1), and BUPS2 form complexes with wall-anchored leucine-rich repeat extensins (LRX8/9/11) in Arabidopsis (*Arabidopsis thaliana*) pollen and bind to RALF4 and RALF19 to sense alterations in the cell wall ^20,21^. In pollen, *Cr*RLK1L-mediated CWI signaling also involves mechanosensitive ion channels, such as MscS-Like8 (MSL8)^22^. Besides CWI, FER also plays essential roles in root-related biological processes, such as the response to soil-borne disease^23^ and monitoring energy and nutrient availability^13,24^. Loss-of-function *FER* mutants (such as the *fer-4* allele) exhibits defects in root hair development^25-27^ and have limited mechanosensitivity, resulting in an abnormal growth response to various mechanical stimuli^28^. In addition, *fer-4* roots acidify their surrounding growth medium faster than the wild type and develop longer roots than the wild type under blue light conditions that stimulate root growth^18^. Moreover, *fer-4* roots are insensitive to the inhibition of root growth triggered by the FER peptide ligand RALF1^18^. FER therefore acts as a regulator hub for root growth via linking external environmental signaling with internal cellular activity^13^. Based on these observations, we were curious about whether and how FER might sense mechanical stimuli to modulate root penetration into the soil.

In this study, we established that the *fer-4* mutant shows defects of root penetration into hard soil and medium. We then conducted single-cell transcriptome deep sequencing (scRNA-seq), which revealed that *fer-4* root cells exhibit a distinct clustering profile compared to the wild type, especially for the cells with root hair and root cap identity. This transcriptome analysis allowed us to also identify several putative FER-regulated transcription factors, one of which being PIF3. A combination of biochemical, imaging and genetic approaches confirmed that FER interacts with, phosphorylates, and stabilizes PIF3, which then modulates the expression of its target genes in the root cap, such as the mechanosensitive ion channel *PIEZO* and genes related to cell sloughing, to confer the capacity to penetrate soil.

## Results

### FER participates in root penetration

We determined whether FER is involved in root penetration, by growing the wild-type Col-0 and the loss-of-function *fer-4* mutant^25^ in soil (low physical barrier) and sand (high physical barrier). Compared to Col-0, *fer-4* developed a shorter primary root (Fig. 1A, B and Exteneded Data Fig. 1A). In addition, *fer-4* seedlings tended to show nonpenetrating roots that instead curved into the air or into the sand (bottom panel in Fig. 1C), likely explaining the weak growth of these mutant seedlings (upper panel in Fig. 1C). Based on these observations, we hypothesized that *fer-4* roots may have difficulty penetrating hard soil. To simulate the growth status of plants in such hard soil conditions, we grew Col-0 and *fer-4* seedlings on the surface of half-strength Murashige and Skoog (MS) medium containing 1%, 1.5%, or 2% (w/v) agar (Fig. 1D). We used the rate of root penetration into the medium as a readout for seedling adaptability to the mechanical constraints imposed by the growth milieu^29-31^. When grown on medium solidified with 1% of agar, most Col-0 and *fer-4* roots were able to grow into the medium (> 95%), with a slightly lower rate for *fer-4* (Fig. 1 D and E). As the agar concentration increased, the rate of root penetration declined in both genotypes (Fig. 1 E), indicating that a harder medium impedes root penetration. Importantly, the penetration rate of *fer-4* roots was more than 20% and 50% lower than that of Col-0 at 1.5% and 2.0% of agar, respectively (Exteneded Data Fig. 1 D and E). In addition, hard medium (2% agar) caused a more severe inhibition of primary root elongation (for medium-penetrating seedling) in *fer-4* compared to Col-0 (Exteneded Data Fig. 1B). These results indicated that *fer-4* roots have a defect in penetrating harder medium. We confirmed this phenotype with a null FER mutant in the C24 accession, *sirène* (*srn*, Exteneded Data Fig. 2A)^25,32^. In addition, a knock-out mutant in LORELEI (LRE)/LORELEI-LIKE-GPI-ANCHORED PROTEIN 1 (LLG1), *llg1-2*, lacking activity of a RALF coreceptor that works with FER^33^, also showed a defect in root penetration into harder medium (Exteneded Data Fig. 2B-2C). We also tested whether FER function in the regulation of root penetration is conserved in other dicotyledons by quantifying the root penetration rate of the two soybean mutant alleles *Gmlmm1-1* and *Gmlmm1-2*, which carry mutations in the FER paralog LESION MIMIC MUTANT1^34,35^. As with *fer-4*, both *Gmlmm1-1* and *Gmlmm1-2* displayed a lower penetration rate and a larger root cap, than their parental cultivar Williams82 (Exteneded Data Fig. 3A-3B). In addition, when growing from uncompacted soil to compacted soil, *Gmlmm1-1* showed a larger bending angle than Williams82, that is θ_2_ > θ_1_ (Exteneded Data Fig. 3C), suggesting a defect of *Gmlmm1-1* in penetrating into the compacted soil. We concluded that FER signaling plays an essential role in root penetration into soil.

**Fig. 1.**
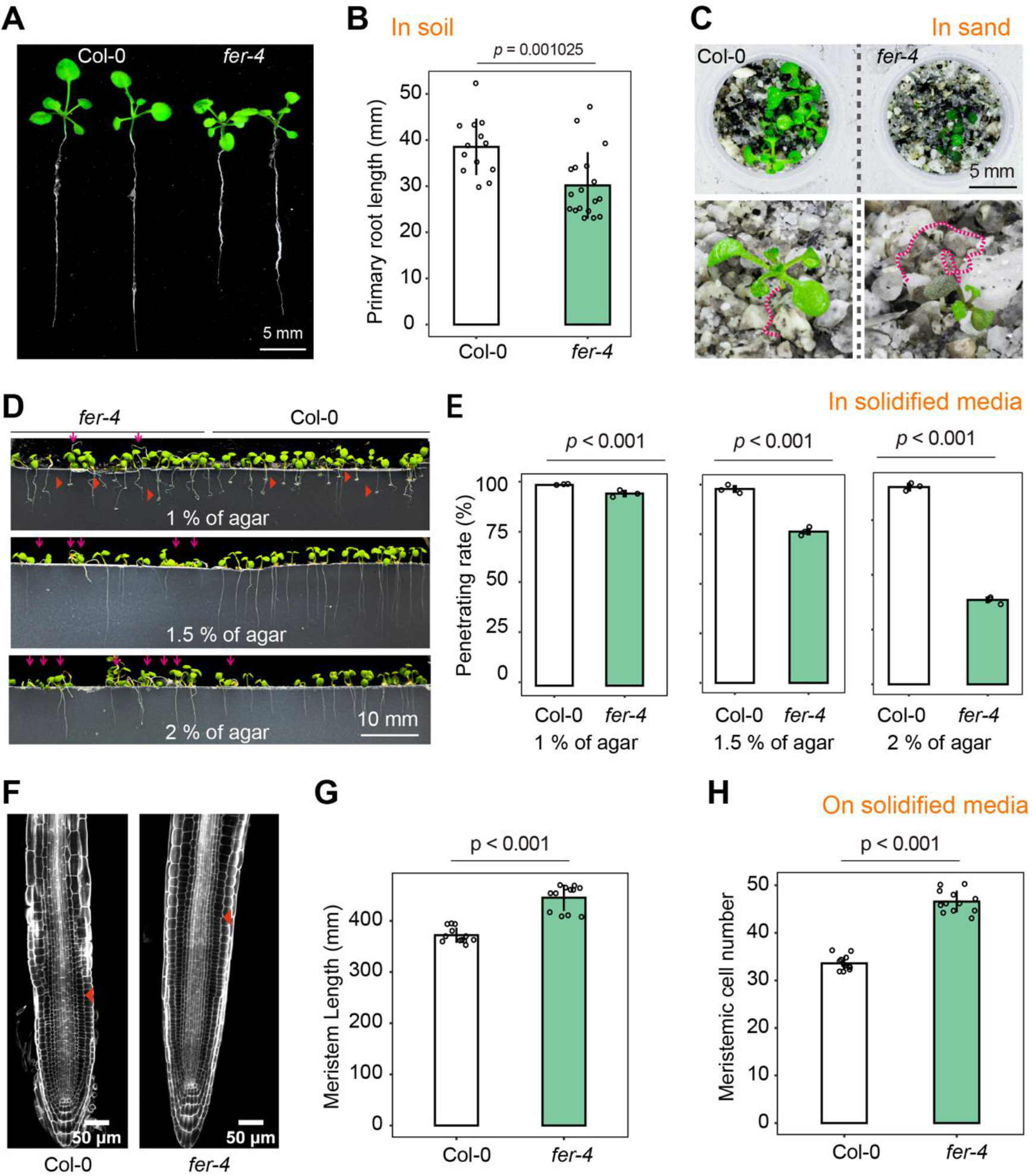
*fer-4* defects in root penetration into soil and root architecture. A-B. Three-week-old Col-0 and *fer-4* seedlings grown in soil. Primary root length (B). C. Representative phenotype of 10-d-old Col-0 and *fer-4* seedlings grown on sand for 10d. The dashed magenta line indicates the root trajectory. D. Representative images showing growth status of primary roots from 7-d-old Col-0 and *fer-4* seedlings grown on agar-solidified medium. The red triangles indicate helical roots. The magenta arrows point to the nonpenetrating root. E. Rate of root penetration of Col-0 and *fer-4* seedlings in growth medium with different agar concentrations. Data are means ± SE (n = 3 plates, >20 seedlings in each group). F. Root tips of 7-d-old Col-0 and *fer-4* seedlings grown on the surface of agar-solidified MS medium. The red triangle indicates the edge of the meristem. G-H. Meristem length (G) and number of root cortical meristematic cells (H) of Col-0 and *fer-4* seedlings shown in F. Data are means ± SD from three independent experiments (n > 15 seedlings). At least three biological replicates in (A)–(H) were performed with similar results. Data are shown as the mean ± SD; for B, E, G and H, *P* values were obtained by Student’s *t*-test comparing Col-0 and *fer-4*.

Root adaptations to mechanical impedance comprise three possible strategies: 1) increased production of mucilage at the root tip and root cap cell sloughing that lubricates the root soil interface^6^; 2) sharper root tip shapes reduce stress at the root tip via a more cylindrical cavity expansion^36,37^; and 3) root hairs help root tip penetration by anchoring the root into the soil^8^. Therefore, the root architecture requires a strict spatiotemporal organization and influences root penetration ability^9^. We thus asked whether the defect in root penetration seen in *fer-4* is associated with its root architecture. *fer-4* seedlings had fewer and shorter root hairs and longer meristems with more cells (Exteneded Data Fig. 1F-H) and a larger root cap (Exteneded Data Fig. 1D-1E, and see below) than those of Col-0. In addition, *fer-4* (Exteneded Data Fig. 1C) and *llg1-2* (Exteneded Data Fig. 2D) roots had a wider cortex and/or root diameter at the quiescent center position. Collectively, *fer-4* showed defects in root penetration and displayed an abnormal root architecture, hinting at a link between these two phenotypes.

### Single-cell RNA-seq analysis of Col-0 and *fer-4* roots

Root architecture and development correlates with root penetration capacity^10,11,38^. To determine the mechanism(s) of FER in root penetration, we performed single-cell transcriptome deep sequencing (scRNA-seq) of *fer-4* and Col-0 root tips. Accordingly, we treated 7-d-old Col-0 and *fer-4* seedlings with water (Mock) or with 2 μM RALF1 (to activate FER) for 2 h, before isolating protoplasts for transcriptome analysis using the 10X Genomics Chromium platform^39^. We detected approximately 2,430 expressed genes per cell, with more than 24,000 total genes across the cell population. Data were prefiltered at both cell and gene levels, resulting in a pool of 26,376 *fer-4* cells and 44,529 Col-0 cells used for further analysis (see Methods). We accounted for the effect of protoplasting on the transcriptome by regressing out the variations caused by protoplasting-induced genes^40^, which were removed from the analysis. After multiple quality control steps (see Supplementary Materials and Exteneded Data Fig. 4A), we conducted a cell-type annotation and cluster identification. First, we uncovered 19 distinct clusters for 12 samples of Col-0 and *fer-4* roots after principal component analysis (PCA) and unsupervised analysis (Exteneded Data Fig. 4B). Importantly, the clustering pattern of *fer-4* cells was distinct from that of Col-0 based on their scRNA-seq profiles (Fig. 2A). Second, we subjected the six samples from Col-0 roots and the six samples from *fer-4* roots to a separate clustering analysis using uniform manifold approximation and projection (UMAP, Exteneded Data Fig. 2B and 2C). Third, after assigning each cluster to their likely cell type based on the expression levels of known marker genes^41,42^ (Exteneded Data Fig. 4C), we observed that nearly all major cell or tissue types present in the root were captured by at least one cluster (Exteneded Data Fig. 2B and 2C). Notably, the *fer-4* samples showed distinct clustering for the cell types of cortex (Cortex_b) and non-hair epidermis (Non-hair epidermis _b) from those obtained with Col-0 samples (Cortex_a and Non-hair epidermis_a, Exteneded Data Fig. 2B and 2C) via UMAP. Moreover, cells from the lateral root cap of *fer-4* (Lateral root cap_a/_b) also showed a clustering pattern different from those of Col-0 root caps (Lateral root cap, Exteneded Data Fig. 2B and 2C), suggesting that FER is involved in the development of these tissues. Collectively, scRNA-seq profiling revealed a developmental role for FER in the root, which is useful for determining the mechanism(s) of FER in root penetration.

**Fig. 2.**
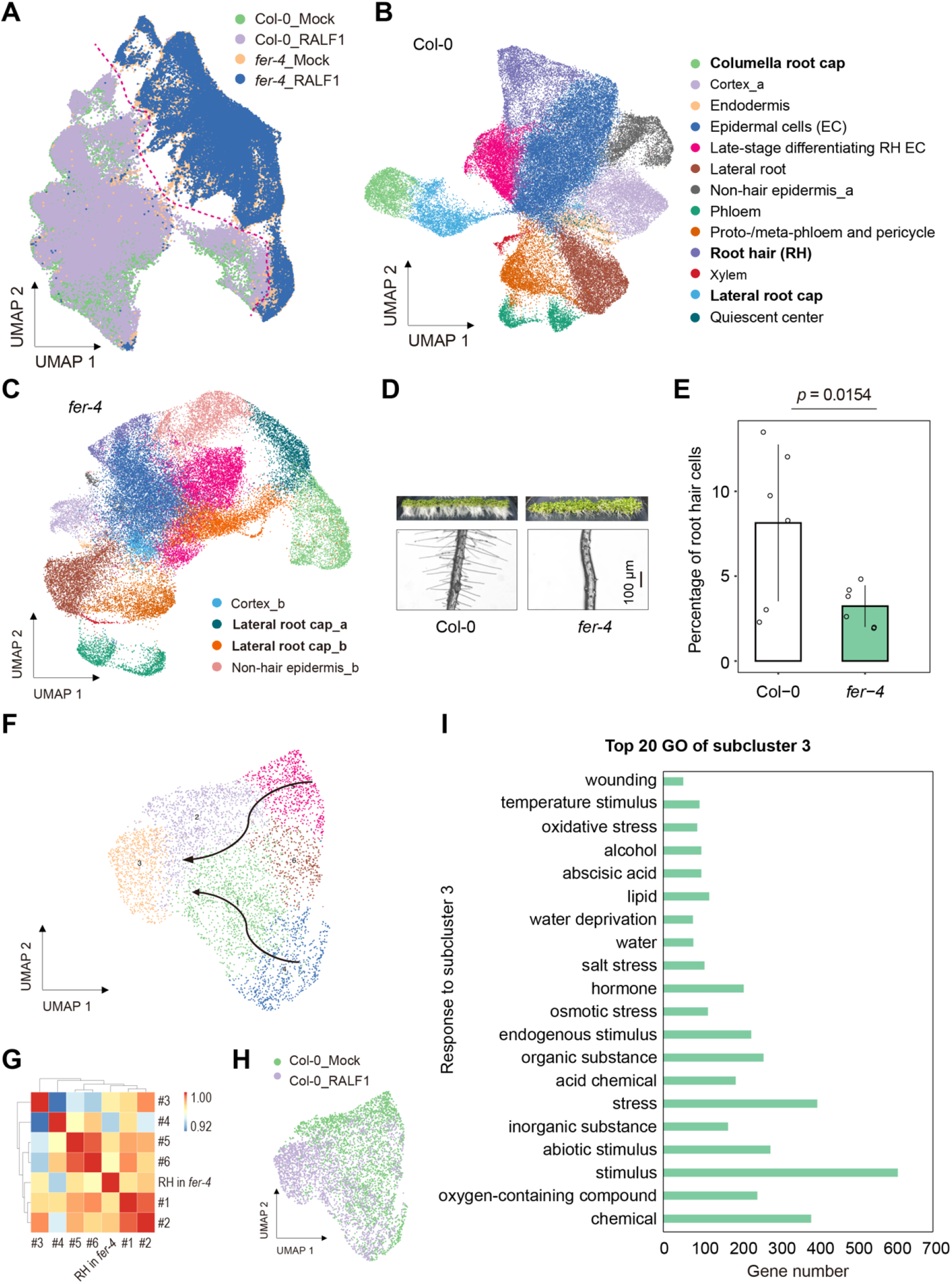
Annotation of cell types from scRNA-seq results of Col-0 and *fer-4* Arabidopsis roots. A. UMAP representation of 12 samples from Col-0 and *fer-4* root cells under mock conditions or treated with RALF1, colored by groups. Colors denote the corresponding cell clusters. B-C. UMAP representation of six Col-0 (B) and six *fer-4* (C) samples. D. Representative phenotypes of root hairs from Col-0 and *fer-4* seedlings grown on half-strength MS agar-solidified medium. E. Percentage of root hair cells in each scRNA-seq sample from Col-0 and *fer-4*. Data are means ± SD. The *P*-value was obtained by Student’s *t*-test. F. Sub-clustering of Col-0 root hair cell type. Arrows indicate developmental trajectory. G. Correlation analysis for root hair cells in *fer-4* and subcluster of root hair in Col-0. H. Subcluster UMAP representation as in B, colored by treatment. I. GO analysis for genes from the RALF1-dependent subcluster (#3). The top 20 terms are shown.

In agreement with the root hair phenotype of *fer-4*, we captured fewer cells with a root hair signature by scRNA-seq (Exteneded Data Fig. 2D and 2E). Root hair development can be divided into three phases: cell specification, initiation of bulge formation, and polarized tip growth^43^. Root hair development is associated with transcription and translation during cell specification, while stress responses and lipid biosynthesis occur at the late stage of polarized tip growth. To characterize root hair development at transcription level, we selected the root hair cell type of Col-0 and subclustered it using UMAP (Fig. 2F). We then used the differentially expressed genes (DEGs) obtained for each subcluster for a Gene Ontology (GO) analysis. We determined that subclusters 4 and 5 represents the cell specification stage, while subcluster 3 matched the late stage (Fig. 2G). We also plotted the six subclusters and extracted pseudotime information, which revealed the developmental trajectory between subclusters, starting in subcluster 4 and 5 and ending in subcluster 3 (Fig. 2F). To determine the stage of the *fer-4* root hairs, we conducted a correlation analysis between the expression profile of *fer-4* root hairs and the six subclusters from Col-0 root hair. Based on scRNA-seq data, *fer-4* root hair appeared closer to subclusters 1 and 2, suggesting that FER is involved in root hair development and that most *fer-4* root hairs can enter the maturation stage without FER. Our previous study showed that RALF1, a ligand of FER, activates the FER signaling pathway to mediate root hair growth^26^. After a 2h RALF1 treatment, the transcriptome of root hairs in Col-0 displayed a clear difference relative to mock-treated root hairs, as evidenced by UMAP clustering (Fig. 2H). GO analysis revealed that RALF1 treatment altered the expression of pathways related to stimulus (Fig. 2I). These findings supported the notion that the RALF-FER pathway mediates root hair growth, which is consistent with previous work^25,26^ and validate our scRNA-seq approach. We thus exploited the scRNA-seq data in more detail to analyze the mechanism(s) by which FER affects root penetration.

### Transcription factors and gene expression patterns regulated by FER in the root cap

As the root cap is a key tissue for root penetration^10,11,38^ and *fer* mutants (*fer-4, Gmlmm1-1* and *Gmlmm1-2*) had more cells in their root caps (for both the columella and lateral root cap, Fig. 3A) relative to the respective wild types (Col-0 and Williams 82, Fig. 3B and 3C, Exteneded Data Fig.3B), we focused on FER-regulated transcription factors (TFs) in this tissue. To identify the primary TFs responsible for cell type-specific DEGs between *fer-4* and Col-0, we predicted cell type-specific TFs regulated by FER using the integrated Gene Regulatory Network (iGRN)^44^ and Motif Enrichment Tool (PMET)^45^. The overlapping TFs defined by these two approaches are listed in Fig. 3D. We identified 8, 15, and 11 overlapping TFs for cell types columella root cap, lateral root cap_a, and lateral root cap _b, respectively (Fig. 3D). Of these TFs, PIF3 and PIF4 were present in all three cell types, while MYC2, MYC3, and MYC4 were present in at least two cell types (Fig. 3D). Notably, MYC2 physically interacts with FER and activates jasmonic acid signaling, thus negatively contributing to plant immunity^46^, in addition to cooperating with FER to regulate touch signaling in aerial tissues^47^. Mutants in *MYC2* or related genes did not show a defect in root penetration into any of the agar-solidified media tested (see below), suggesting that MYCs may have a negligible contribution. We thus focused on the roles of PIF3 in regulating gene expression in the root cap.

**Fig. 3.**
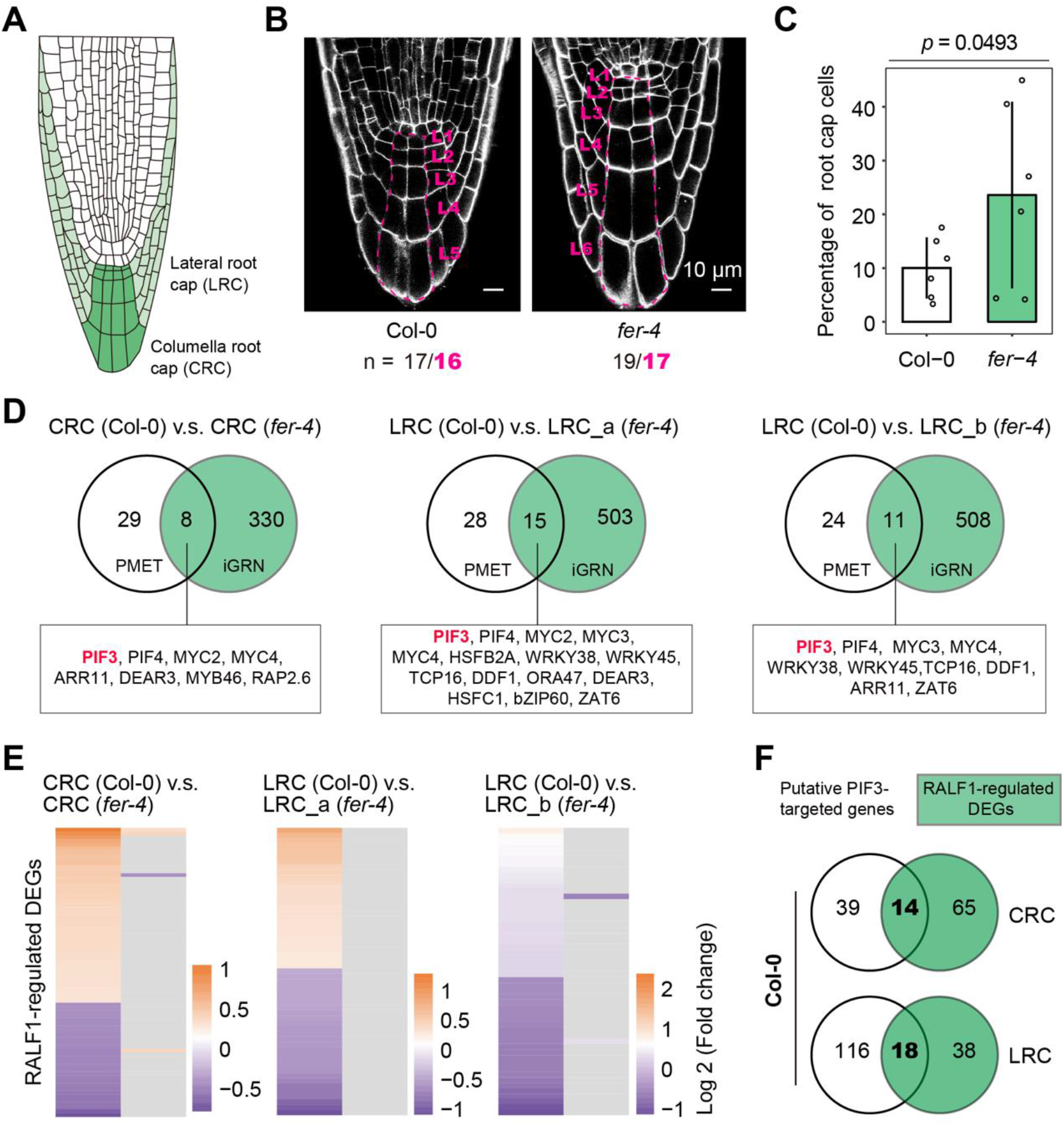
Analysis of transcription factors that are regulated by FER. A. Schematic diagram of the Arabidopsis root cap. B. Columella root cap in Col-0 and *fer-4*. C. Percentage of root hair cells in each scRNA-seq sample from Col-0 and *fer-4*. Data are means ± SD. The *P*-value was obtained by Student’s t-test. D. Prediction of transcription factors regulated by FER in the columella and lateral root caps using PMET and iGRN approaches. E. RALF1-induced differentially expressed genes between Col-0 and *fer-4* in root cap cells. F. Venn diagram showing the number of overlapping DEGs between putative PIF3-target genes and RALF1-regulated genes. CRC: columella root cap, LRC: lateral root cap.

We investigated whether RALF1 affects gene expression in the root cap via PIF3, by identifying DEGs whose expression changed in response to RALF1treatment in root cap cells of Col-0 and *fer-4* (Fig. 3E). In both CRC and LRC, we observed that dozens of genes are regulated by RALF1 treatment in Col-0, while the same genes showed no change in expression in *fer-4* (Fig. 3E), which was consistent with FER acting as a *bona fide* RALF1 receptor^18^. A comparison of RALF1-regulated DEGs to known PIF3-target genes (DEGs used to enrich for PIF3 in iGRN), yielded 14 (26.4% for PIF3-target genes) and 18 (13.4%) common DEGs in CRC and LRC, respectively (Fig. 3F). This result indicated that PIF3 is an important TF that modulates gene expression in the root cap.

### PIF3 regulates root penetration through mediating root cap sloughing and the expression of mechanosensitive genes

To ascertain the potential TFs acting downstream of FER in the context of root penetration, we determined the penetration capacity of their corresponding mutants (Exteneded Data Fig.7). With the exception of *pif3* and *wrky38* mutants that displayed a lower penetration rate than Col-0 when grown on 2% agar medium, the mutants *arr11* (*Arabidopsis response regulator 11*), *rap2*.*6* (*related to ap2*.*6*), *wrky45, dff2* (*dwarf and delayed flowering*), and *zat6* (*zinc finger of arabidopsis thaliana 6*) as well as the higher-order *myc* mutants *myc2myc3, myc2myc4, myc3myc4* and *myc2myc3myc4* showed penetration rates comparable to that of Col-0 in 1% to 2% agar medium (Fig. 4A and Exteneded Data Fig.7A). Importantly, *pif3-1* and *pif3-3* roots showed a lower penetration rate in 2% agar medium than Col-0 (Fig. 4A) that was reminiscent of *fer-4*. In addition, the CRC in *pif3-1* and *pif3-3*, like *fer-4*, had one additional cell layer compared to Col-0 (Fig. 4B and C). A line overexpressing PIF3, *35S::PIF3-myc* (*PIF3-myc*), displayed a normal penetration rate and cell layer in the CRC. Unlike *pif3* mutants, *wrky38* showed a cell layer number in the CRC comparable to that of Col-0 (Exteneded Data Fig. 7B). Collectively, these data indicated that PIF3 is also required for root penetration.

**Fig. 4.**
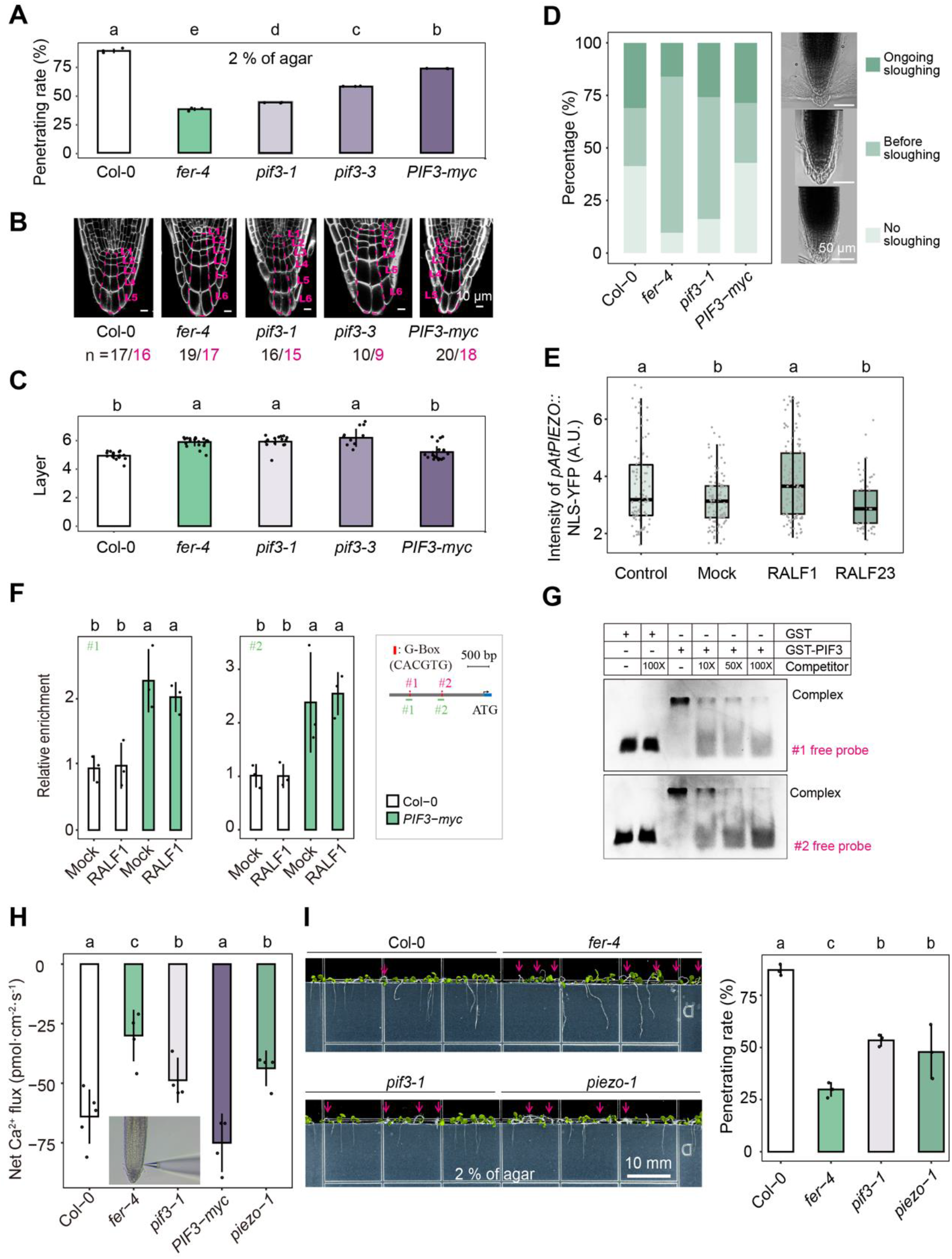
PIF3 regulates root penetration through mediating root cap sloughing and the expression of a mechanosensor gene. A. Rates of root penetration by Col-0, *fer-4, pif3-1, pif3-3*, and *PIF3-myc in Col-0* (*PIF3-myc*) into medium solidified with 2% agar. Data are presented as mean ± SE (n = 3 plates, >20 seedlings of each group). B. Root cap phenotype of Col-0, *fer-4, pif3-1, pif3-3*, and *PIF3-myc*. The dashed magenta line indicates the columella root cap (CRC) cells. L: layer. C. Cell layer number of in the CRC in *fer-4, pif3-1, pif3-3*, and *PIF3-myc*. D. Root cap sloughing analysis for Col-0, *fer-4, pif3-1*, and *PIF3-myc*. More than 80 roots were counted for each group. E. AtPIEZO-NLS-YFP fluorescence intensity in root cap cells after treatment with RALFs. F. ChIP-qPCR analysis of PIF3 binding at the *PIEZO* promoter. ChIP assays were performed using an anti-Myc antibody. Data are means of three biological replicates ± SD. G. PIF3 bound to two motifs within the *PIEZO* promoter. Competitor was unlabeled DNA fragments. Probes used for the EMSA are shown in F in magenta. H. The net Ca^2+^ fluxes in Col-0, *fer-4, pif3-1, PIF3-myc* and *piezo-1* seedlings that were grown on the medium for 3d. The results were analyzed with one-way ANOVA. Different lowercase letters indicate significant differences at *P* < 0.05. The inset diagram indicates the position for Ca^2+^ flux measurement using NMT. I. Rates of root penetration of Col-0, *fer-4, pif3-1*, and *piezo-1* seedlings into medium solidified with 2% agar. Data are means ± SE (n = 3 plates, >20 seedlings of each group). Magenta arrows indicate nonpenetrating roots. For (A, C, E, Fand H), data were analyzed by one-way ANOVA with Tukey’s test. Different lowercase letters indicate significant difference (*P* < 0.05).

Root cap sloughing helps protect roots when they penetrate the soil^48,49^. Higher levels of pectin methyl esterification of the cell wall attenuate the sloughing of the root cap outer layer by promoting cell-cell adhesion^50^. PIF3 was shown to regulate the expression of *Pectin Methylesterase Inhibitors* genes (*PMEIs*, At5g62330, At5g62360 [PMEI13], At4g25260 [PMEI7], At5g04970)^51^, which determine pectin methyl esterification levels in the cell wall. We therefore characterized the sloughing phenotypes of the root cap from Col-0, *fer-4, pif3-1*, and *PIF3-myc* seedlings. We defined three phases to describe the state of root cap sloughing: no sloughing, before sloughing, and ongoing sloughing. More root caps of *fer-4* and *pif3-1* mutant seedlings were in the before sloughing phase than Col-0 (Fig. 4D), suggesting that their root cap outer layer is defective in sloughing. In contrast to *fer-4* and *pif3-1, PIF3-myc* and Col-0 seedlings had a similar distribution for each phase of root cap sloughing (Fig. 4D). This result suggested one strategy whereby PIF3 controls root cap sloughing by regulating the expression of cell wall remodeling genes.

Next, we asked whether the FER-PIF3 module regulates the expression of mechanosensitive genes. PIEZO was previously reported to be expressed in the root cap and is required for root penetration into hard medium^29,31,52^, prompting us to measure its expression levels in response to RALF-mediated FER activation. For this purpose, we used a transgenic line expressing a construct encoding a nucleus-targeted yellow fluorescent protein (YFP) under the control of the Arabidopsis *PIEZO* promoter (*pAtPIEZO::NLS-YFP*)^29^. We detected elevated YFP fluorescence in root cap cells upon treatment with RALF1, but not with RALF23 (Fig. 4E). We also performed chromatin immunoprecipitation (ChIP) assays to assess the targeting of *PIEZO* by PIF3, using Col-0 as a negative control and *PIF3-myc* (in Col-0 background) lines as a test sample. We established that PIF3 is enriched over *PIEZO* chromatin regions containing the G-box (CACGTG) elements in *PIF3-myc* but not in Col-0, suggesting that PIF3 binds to the *PIEZO* promoter *in vivo* (Fig. 4F). We confirmed this result by electrophoretic mobility shift assay (EMSA) using recombinant glutathione S-transferase (GST)-tagged PIF3 (Fig. 4G). Consistently, *PIEZO* transcript levels were lower in the roots of *pif3-1* and *fer-4* plants than in those of the wild type, whereas transgenic expression of *PIF3-myc* rescued this defect (Fig. S8). These results suggest that PIF3 directly binds to the *PIEZO* promoter and regulates its transcription.

*PIEZO* changes the Ca^2+^ flux in the root cap^29^. Similar to *piezo-1*, both *fer-4* and *pif3-1* displayed weaker net Ca^2+^ fluxes in the root cap in the non-damaging micro-measurement technique (NMT) assay, whereas *PIF3-myc* and Col-0 root caps had a similar net Ca^2+^ flux (Fig. 4H). In line with the above results, *piezo-1* roots exhibited a penetration defect similar to that seen in *pif3-1* and *fer-4* (Fig. 4I). These findings indicate that the FER-PIF3 module regulates *PIEZO* expression in the root cap.

### FER interacts with, phosphorylates, and stabilizes PIF3

To further explore the molecular mechanisms by which the FER pathway modulates root penetration, we tested whether FER interacts with PIF3 in yeast two-hybrid (Y2H) assay. Indeed, we observed that PIF3 interacts with the cytoplasmic domain of FER, FER-CD (amino acids 469-896, Fig. 5A)^53,54^. Co-immunoprecipitation (Co-IP) assays with recombinant PIF3-GST or genetically crossed *FER-GFP* and *PIF3-myc* plants (Fig. 5B) indicated that FER associates with PIF3 *in vitro* and *in vivo* ((Fig. 5B, Exteneded Data Fig. 9). In agreement with the Y2H and Co-IP results, pull-down experiments also suggested that PIF3-GST can interact with His-tagged kinase domain of FER (518-816 amino acids, FER-KD) *in vitro* (Fig. 5C). As the interaction between the kinase and PIF3 raised the possibility that FER might phosphorylate the TF, we performed *in vitro* phosphorylation assays with purified recombinant PIF3-GST, His-FER-KD, and His-FER^K565R^-KD (a kinase-dead FER, as negative control). As shown in Fig. 5D, an antibody against phosphorylated serine residues (anti-pSer/Thr) detected recombinant PIF3 when PIF3-GST was co-incubated with His-FER-KD (lane 3), but not with the negative control His-FERK565R-KD (lane 5). His-FER-KD was detected because of its autophosphorylation ability^55,56^. We identified 10 phosphorylation sites (Ser-48, Ser-58, Ser-115, Ser-160, Ser-196, Ser-201, Thr-234, Ser-250, Thr-335, and Thr-499) in this phosphorylation system using LC-MS/MS (Fig. 5E and Exteneded Data Fig.10). Notably, 8 of these 10 phosphorylation sites (Ser48, Ser-115, Ser-160, Ser-196, Ser-201, Thr-234, Thr-335, and Thr-499) were distinct from those known to be targeted by the red light photoreceptor phytochrome, phyB^57^, suggesting that FER-dependent phosphorylation of PIF3 may exert a function different from that of phyB. To test whether these phosphorylation sites are required for PIF3 function, we generated transgenic lines in *pif3-1* background that expressed the phospho-mimicking (*pPIF3::PIF3*^*mut10D*^*-Venus*, 10 phosphorylation sites mutated to Asp), phospho-dead (*pPIF3::PIF3*^*mut10A*^*-Venus*, 10 phosphorylation sites mutated to Ala), and wild-type PIF3 variants under control of PIF3 promoter (Exteneded Data Fig.11A). Although *PIF3* transcript levels are comparable between *pPIF3::PIF3-Venus, pPIF3::PIF3*^*mut10A*^*-Venus* and *pPIF3::PIF3*^*mut10D*^*-Venus* (Exteneded Data Fig.11B) and the phenotype of root soil penetration was recovered in *pPIF3::PIF3-Venus* and *pPIF3::PIF3*^*mut10D*^*-Venus* (Exteneded Data Fig.11C), we failed in screening of a line of *pPIF3::PIF3-Venus* and *pPIF3::PIF3*^*mut10A*^*-Venus* with strong fluorescence intensity of Venus as pPIF3::PIF3^mut10D^-Venus, *Empty vector-Venus* or *PIF3-GFP* (driven by 35S promoter, Exteneded Data Fig.11A). To circumvent this difficulty, we performed *in vivo* stability test in *PIF3-GFP*. Activation of FER upon RALF1 treatment increased PIF3 abundance (Fig. 5F) and its accumulation in the nucleus of root cells (Fig. 5G). To determine whether phosphorylation by FER stabilizes PIF3, we tested the degradation rate of recombinant PIF3 and its phospho-mimicking (GST-PIF3^mut10D^) and phospho-dead (GST-PIF3^mut10A^) versions by the lysate of Col-0 roots within 5 min. GST-PIF3^mut10D^ displayed a lower degradation that GST-PIF3 and GST-PIF3^mut10A^ (Fig. 5H), indicating that the phosphorylated PIF3 is more stable. This may explain the fact that *pPIF3::PIF3*^*mut10D*^*-Venus*, rather than *pPIF3::PIF3-Venus* or *pPIF3::PIF3*^*mut10A*^*-Venus*, displayed a observable signal of Venus in root cells (Exteneded Data Fig.11A). Given that the binding of PIF3 to the *PIEZO* promoter was not affected by RALF1 treatment (Fig 4F), we speculated that PIF3 stabilization mainly contributes to facilitate the expression of *PIEZO* in the root cap (Fig 4E). Lastly, overexpression of PIF3 in the *fer-4* background (*fer-4 PIF3-myc*) partially rescued the defect in penetration capacity of the *fer-4* mutant when grown on MS medium solidified with 1.5% and 2.0% agar (Fig. 5I) and partially restored CRC phenotype (Exteneded Data Fig.12). Also, overexpression of PIF3 promoted the growth of primary root, hypocotyl, petiole, and shoot of *fer-4* (Exteneded Data Fig.13). Mutation of *PIF3* did not show those effects (Exteneded Data Fig.12 and 13). Collectively, these data indicate that FER physically interacts with, phosphorylates, and stabilizes PIF3.

**Fig. 5.**
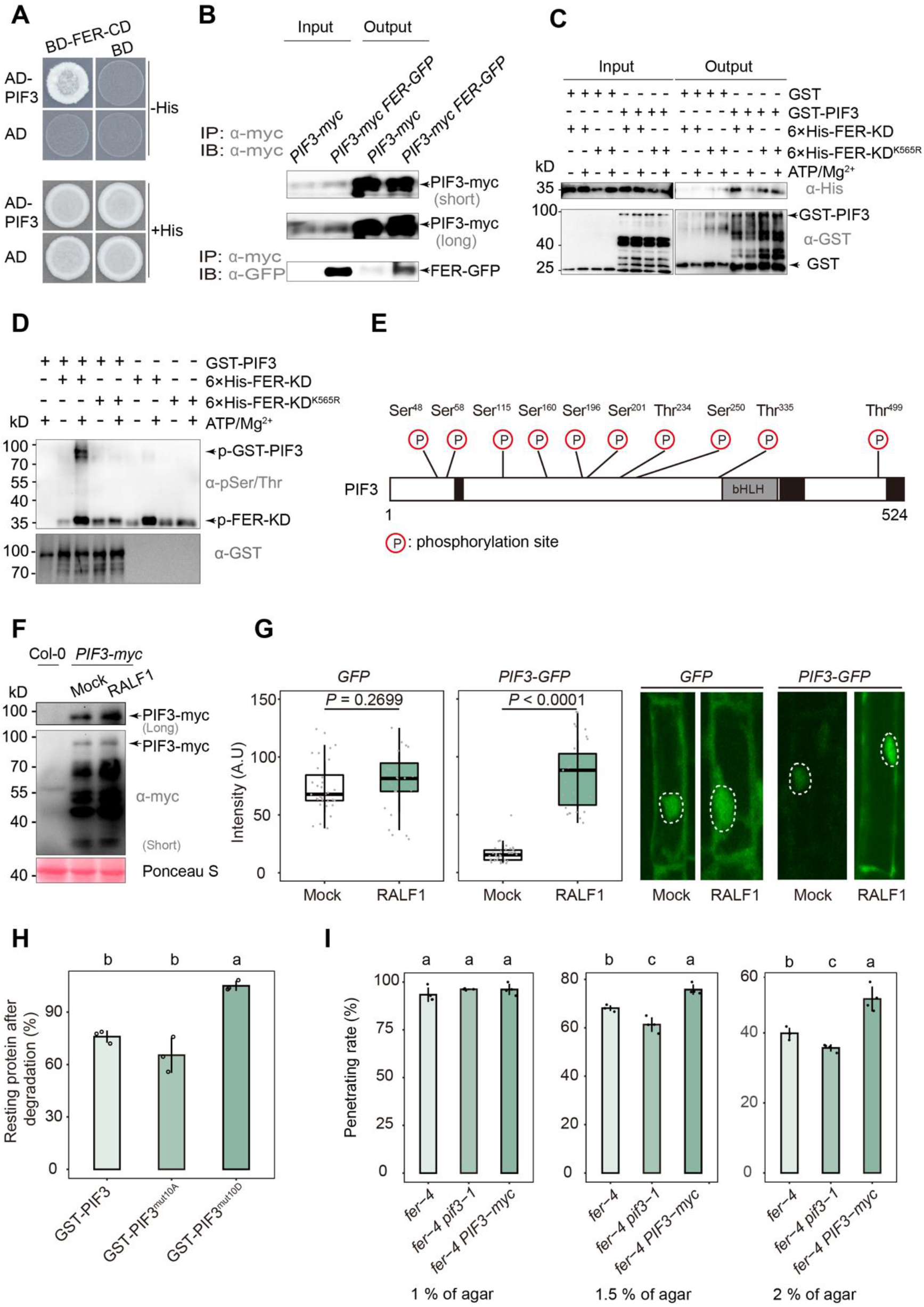
FER interacts with, phosphorylates, and stabilizes PIF3. A. Y2H assays showing the interaction between PIF3 and FER. Synthetic dropout medium (His) containing 20 mM 3-amino-1,2,4-triazole (3-AT) was used to test the interaction. PIF3 was cloned into the pGADT7 (AD) vector, and the FER-CD was cloned into the pGBKT7 (BD) vector. B. Co-IP assays with recombinant PIF3-GST and FER-GFP, using an anti-GST antibody for the IP. PIF3-GST and FER-GFP were detected using anti-GST and anti-GFP antibodies, respectively. C. GST pull-down assay. The GST protein and His-FER-KD proteins were detected with anti-GST and anti-His antibodies, respectively. D. *In vitro* phosphorylation assay for PIF3-GST. E. Schematic diagram of PIF3 with all phosphorylation sites identified in PIF3. F. PIF3-myc abundance increases upon treatment with RALF1 for 5 h. G. Nucleic GFP fluorescence intensity of the root cells of *GFP* and *PIF3-GFP* seedlings after treatment with RALF1 for 2h. Boxplots span the first to the third quartiles of the data. The *P*-values were obtained by Student’s *t*-test. The nuclei are highlighted by the dashed white circles. H. PIF3-related recombinant proteins in the *in vitro* degradation assay. The abundance of GST-tagged protein was determined by in immunoblot analysis. Different lowercase letters indicate significant differences at *P* < 0.05. I. Behavior of *fer-4, fer-4pif3-1* and *fer-4 PIF3-myc* seedlings in the penetration assay. Data were analyzed by one-way ANOVA with Tukey’s test. Different lowercase letters indicate significant differences (P < 0.05)

## Discussion

In nature, plant roots avoid physical impedance to penetrate into soil to obtain water and nutrients. Here, we propose a signaling pathway whereby the CWI sensor FER works in concert with the mechanosensitive ion channel PIEZO (via PIF3) in roots to perceive soil constraints to modulate soil penetration capacity. Our findings also indicated that the basic helix-loop-helix (bHLH) TF PIF3 unifies two key processes of seed germination: cotyledon emergence above the soil surface^1-3^ and root penetration into the soil (this work, Fig. 6).

**Fig. 6.**
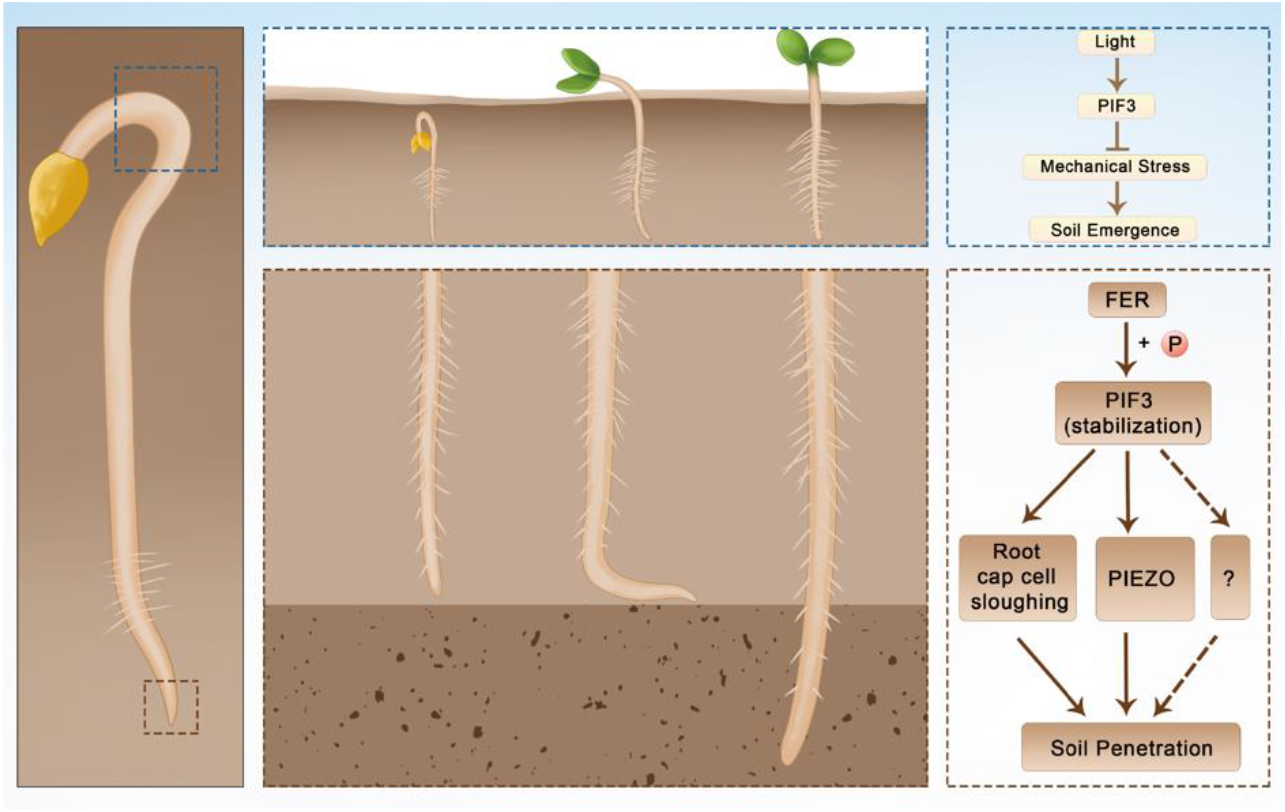
A working model for the integrated regulation of the emergence of cotyledons from the soil and the penetration of the root into the soil by PIF3. For the cotyledon emergence, PIF3 transduces light signals to control mechanical stress caused by soil covering. For the root penetration, PIF3 transfers the soil constraints perceived by FER to the transcriptional machinery to mediate root cap cell sloughing and induce the mechanosensitive ion channel gene *PIEZO* in the root cap, and possible unknown factor(s). In this pathway, FER phosphorylates and stabilizes PIF3, resulting in its higher abundance in root cap.

Both the emergence from and penetration into the soil directly determine nascent seedling growth. Before reaching the soil surface, seedlings prioritize hypocotyl elongation by monitoring the quantity and quality of light via phytochromes. Whether or not light signaling components are involved in the regulation of root growth into the deeper soil remains unexplored. We identified a receptor kinase-transcription factor module that integrates mechanical pressure and darkness signals: In the absence of light, PIF3, probably with a photoreceptor such as phototropin-1^58^, transduces the mechanical signal perceived by FER to the transcriptional level to confer root penetration capacity to Arabidopsis seedlings. Although FER was shown to perceive mechanical signals in seedlings^28^, the site perception was missing. The present study also demonstrates that FER senses soil constraints through the root cap, leading to the regulation of cell wall remodeling and the mechanosensitive ion channel genes. Importantly, our findings do not rule out the possibility that root hairs and the cortex might contribute to root penetration as well. Based on our scRNA-seq profiling, their roles can be further explored. Given that FER functions in nutrient balancing^24^, osmotic stress^15^, light signaling^58^, and plant-bacteria interactions^59^, future work should focus on determining whether the FER-mediated penetration pathway steers the direction and depth of root growth via these related pathways.

## Supporting information

Combined Supplementary Materials

