## Supplementary material for "The soil emergence-related transcription factor PIF3 regulates root penetration by interacting with the receptor kinase FER": Combined Supplementary Materials

#### Materials and Methods

#### Acknowledgements

#### References

#### Extended Data Fig. 1 to 13

### Materials and Methods

#### 1. Root penetration assay in MS medium

Forty milliliters of half-strength Murashige and Skoog (MS) medium (Murashige and Skoog basal salt mixture [PhytoTechnologies Laboratories]) containing 1%, 1.5% or 2% (w/v) of phytoagar was poured onto a 100 × 100 × 15-mm plate. After it had solidified, the MS medium was cut along a straight line with a sharp sterile blade about 2 cm from one plate edge, and the smaller segment was removed. Arabidopsis *(Arabidopsis thaliana)* seeds were surface-sterilized with 15% (v/v) NaClO for 5 min and then stratified for 2 d at 4 °C in darkness to break dormancy before sowing. The plates were placed vertically at a constant temperature of 22 °C under a long-day (16 h light/8 h dark) photoperiod for 7 d. Seeds were placed on the cut surface at the middle of section (2 mm in thickness). Seedlings whose roots grew into the medium were counted as penetrating and, otherwise as nonpenetrating.

#### 2. *Arabidopsis* seedling growth on sand

River sand (Xiangjiang, Changsha, Hunan) (diameter 250 µm - 1 mm (> 98%), 1 - 3 mm (< 2%)) was sterilized and mixed with Agra-perlite (grain 2.0-3.0 mm) at a 9:1 ratio and placed in round Petri dishes (125 mm in diameter). The sand was sterilized at 120°C for 5h before being pre-soaked with sterilized liquid half-strength MS medium. Seven-day-old seedlings grown on half-strength MS solidified medium were transferred to sand plates (containing a 2-cm layer of sand) and allowed to grow for another 7 d. The growth conditions were the same as above.

#### 3. Calcofluor white staining and confocal microscope imaging of root tips

*In vitro* grown seedlings were fixed overnight in a fixation solution (methanol [50%, v/v], acetic acid [50%, v/v]). Seedlings were then incubated in hot ethanol for 15 min (80% [v/v] ethanol at 80°C) and rehydrated in 50% [v/v] and then 30% [v/v] ethanol for 10 min each. Seedlings were rinsed in distilled water and then incubated in 0.2 M sodium hydroxide containing 1% (w/v) SDS for 30 min at room temperature. Samples were then rinsed in water and stained for 30 min in 1% (v/v) calcofluor white (Fluorescent Brightener 28) to which a few drops of 10 M sodium hydroxide were added to obtain complete dissolution. Seedlings were then rinsed in water and gently mounted on a slide very to avoid squashing the root tips. Samples were excited at 405 nm with an emission band of 410-550 nm with a Zeiss LSM880 confocal microscope using a 40× objective and a GaAsP spectral detector.

#### 4. Soybean soil penetration assay

The soybean cultivar Williams 82, and the *Gmlmm1-1* and *Gmlmm1-2* mutants were a kind gift from Daolong Dou [34]. Soybean seeds were placed on uncompacted soil (1 g $cm^{-3}$) and watered once a day. Seeds were germinated at 28°C before being moved to uncompacted (1 g $cm^{-3}$) and compacted (1.8 g cm$^{-3}$) soil. A layer of uncompacted soil (~5 mm height) was placed on the soil surface to cover the seeds. After 5 d of growth, the soil around the seedlings was removed gently and root penetration was scored. Concerning X-ray computed tomography imaging, seedlings grew in soil containing the uncompacted (1 g cm$^{-3}$) and compacted (1.4 g cm$^{-3}$) layers. The roots of 7-day old soybean seedlings were imaged non-destructively in the soil using Xredia 515 Versa (Carl Zeiss. Inc.) at Analytical Testing Center, Hunan University. Scans were acquired at 110 kV X-ray energy in Normal mode (Objector = 0.4 X, exposure time = 3s, Filter = Air, bin = 2). Scan resolution was 50 μm. Three-dimensional image reconstruction was performed using Scout-and-Scan™ Control System software. Dragonfly V4.2 was used to segment the roots from the soil.

5.Protoplasting

Seeds for the Arabidopsis accession Columbia-0 (Col-0) and the *fer-4* mutant were surface sterilized with 15% (v/v) NaClO, sown on agar plates containing solidified half-strength MS medium (pH 5.8, 0.8% [w/v] sucrose, 1% phytoagar), and grown vertically at 22°C in long-day conditions (16 h light/8 h dark). The roots of 7-d-old seedlings were treated with liquid half-strength MS medium (pH 5.8, 0.8% [w/v] sucrose) containing 2 μM synthesized RALF1 peptide for 2 h. To expose the roots to treatment and to avoid light, a piece of filter paper (1.5 × 8 cm) was soaked in liquid medium containing RALF1 and placed on top of the roots. Approximately 3,000 treated roots were excised approximately 1.5 cm from their tip, broadly sliced with a scalpel, and then passed through a 70-μm cell strainer above a 35-mm well of a six-well plate containing 8% [w/v] mannitol. After rinsing out the debris of solidified medium, the root fragments were treated with 10 mL protoplasting solution optimized for scRNA-seq from a previously described protocol[60]. Immediately before use, the enzyme solution was prepared (1.25% [w/v] cellulase [ONOZUKA R-10, Yakult] 0.1% [w/v] pectolyase [P-3026, Sigma-Aldrich], 8% [w/v] mannitol, 20 mM MES [pH 5.7], 20 mM KCl, 10 mM $CaCl_2$, and 0.1% [w/v] bovine serum albumin). The enzyme solution was activated by heating at 55°C for 10 min. The plate was shaken at 90 rpm for 1 h at 25°C, and then the cell solution was centrifuged at 300 g for 10 min 25°C and the pellet was resuspended in 500 μL washing solution (8% [w/v] Mannitol, 20 mM MES [pH 5.7], 20 mM KCl, 10 mM $CaCl_2$, and 0.1% [w/v] bovine serum albumin). The cell suspension was centrifuged again at 100g for 6 min at 25°C, the entire procedure was repeated twice. All resuspended protoplasts were filtered through a 40-μm cell strainer. The supernatant was discarded and the cells collected by the cell strainer were resuspended with 100 μL of 8% (w/v) mannitol. Eighteen microliters of resuspended protoplasts were mixed with 2 μL of 0.4% (w/v) trypan blue. The stained protoplasts were observed under a microscope, and four photographs were taken for each sample. The living and stained protoplasts and debris were counted and the protoplast density, cell activity, and debris rate were calculated. The cell number was adjusted to ~1,000 cell/μL as recommended by the Chromium 10× Genomics manual. The protoplast suspension was loaded into a Chromium microfluidic chip with 30 v2 chemistry and barcoded with a 103 Chromium Controller (10× Genomics).

8. Single-cell RNA-seq data preprocessing

The Cell Ranger software pipeline (version 3.1.0) provided by 10× Genomics was utilized to demultiplex cellular barcodes, map reads to the Arabidopsis genome BSgenome object ("BSgenome. Athaliana.

TAIR. TAIR9") with TAIR10 gene annotation file using the STAR aligner, and down-sample reads as required to generate normalized aggregate data across samples, producing a matrix of gene counts for each cell. We processed the unique molecular identifier (UMI) count matrix using the R package Seurat version 3.1.1[61]. To remove low-quality cells and likely multiple captures, which is a major concern in microdroplet-based experiments, we removed any cell whose UMI/gene number fell within mean ± 2 standard deviations, assuming a Gaussian distribution of UMI/gene numbers for each cell. Following visual inspection of the distribution of mitochondrial genes expressed in each cell, we further discarded low-quality cells where >10 % of the counts belonged to mitochondrial genes. After applying these quality control criteria, approximately 13,000 single cells were retained for downstream analyses. Library size normalization was performed with the NormalizeData function in Seurat [61] to obtain normalized counts. Specifically, the global-scaling normalization method "LogNormalize" normalized the gene expression measurements for each cell by the total expression levels, multiplied by a scaling factor (10,000 by default), and the results were log transformed.

The top variable genes across single cells were identified using the method described previously[62]. The most variable genes were selected using the FindVariableGenes function (mean.function = ExpMean, dispersion. Function = LogVMR) in Seurat [61]. To remove the batch effects in single-cell RNA-sequencing data, the mutual nearest neighbor method[63] was applied with the R package batchelor. Graph-based clustering was performed to group cells according to their gene expression profiles using the FindCluster function in Seurat[61]. Cells were visualized using 2D following dimensionality reduction according to their gene expression profile using the FindClusters function in Seurat[61] with t-Distributed Stochastic Neighbor Embedding (t-SNE) and Uniform Manifold Approximation and Projection (UMAP) algorithm with the RunTSNE and RunUMAP functions in Seurat, respectively. We used the FindAllMarkers function (test. use = bimod) in Seurat[61] to identify marker genes for each cluster. For a given cluster, FindAllMarkers identified positive markers[64] compared to all other cells.

Differentially expressed genes (DEGs) were identified using the FindMarkers function (test.use = MAST) in Seurat[61]. $P$ value < 0.05 and $|\mathrm{Log_2[foldchange]}| > 0.58$ were set as the thresholds for significant differential expression. Gene Ontology (GO) enrichment and Kyoto Encyclopedia of Genes and Genomes (KEGG) pathway enrichment analysis of DEGs were performed using R, based on the hypergeometric distribution.

7. GO analysis

GO enrichment analysis was performed using Metascape (https://metascape.org/gp/index.html#/main/step1) and the R package ClusterProfiler (version 4.0.5, parameters default). DEGs of the columella root cap, lateral root cap_a, lateral root cap_b, and root hairs were used in Metascape to identify enriched GO terms. The same DEGs were used in ClusterProfiler to select the genes regulated by MYC (MYC2, MYC3, and MYC4) and PIF (PIF3, PIF4, PIF5, and PIF7) transcription factor families. The threshold adjusted $P$ value was < 0.05. All genes in the Arabidopsis genome were used as background. Fisher's exact test was used, and the false discovery rate (FDR) was calculated for multiple test correction. Enriched GO terms were determined for biological processes, molecular functions, and cellular components.

8. Correlation analysis

For correlation analysis of merged single cell and bulk RNA-seq data, $Log_2$ (mean RPM+1) expression values for each gene from two replicates of pooled scRNA-seq and bulk RNA-seq were quantile-normalized, and the Pearson's-correlation coefficient was calculated in R. For correlation analysis between scRNA-seq replicates, each replicate was simulated as a bulk RNA-seq sample, the $Log_2$ (mean RPM+1) expression values were calculated for all genes, and the Pearson's-correlation coefficient between replicates was calculated in R. For correlation analysis between the scRNA-seq replicates across individual clusters, the average expression of cells within a cluster was calculated for each replicate using the Seurat command Average Expression (object, use.raw = T). The Pearson's-correlation coefficient between the replicates was then determined for each cluster using Seurat CellPlot.

9. Paired Motif Enrichment Tool analysis

The identification of pairs of transcription factor (TF) binding motifs was performed with the Paired Motif Enrichment Tool (PMET) (http://nero.wsbc.warwick.ac.uk/tools/user_case_form.php). The gene IDs from TAIR of the above-mentioned DEGs in each cell type were uploaded to the online tool. Parameters were as follows: promoter length = 1,000, max motif matches = 5, number of selected promoters/intervals = 5,000, 5' UTR included, potential overlapped promoter removed. All significant motif pairings with adjusted *P*–value (Bonferroni) < 0.05 were chosen.

10. Integrated Gene Regulatory Network

TF genes associated with DEGs of each cell type were analyzed at http://bioinformatics.psb.ugent.be/webtools/iGRN/ with a set *q*–value < 0.001. The overlapping TFs between iGRN and PMET were identified with TBtools (V 1.0983).

11. Fluorescence intensity measurement

Concerning the expression of *PIEZO* in the root cap, five-day-old Arabidopsis *pAtPiezo::NLS-YFP* seedlings grown on half-strength solidified MS medium were moved to half-strength liquid MS medium containing 1 μM synthesized RALF1 or RALF23 peptides. Seedlings were incubated at a constant temperature of 22 °C for 5 h. For the PIF3 stability test in root cells, 7-d-old *GFP*[56] and *PIF3-GFP*[57] seedlings grown on half-strength solidified MS medium were moved to half-strength liquid MS medium containing 1 μM synthesized RALF1 peptide. Seedlings were incubated in the dark for 2 h. The epidermal cells were observed with an excitation at 488 nm and an emission band of 500-530 nm with a Zeiss LSM880 confocal microscope using a 40× objective and a GaAsP spectral detector. To quantify the relative fluorescence intensity of YFP or GFP, all images were captured using the same laser, pinhole, and gain settings of the confocal microscope to allow comparison of the data. The quantification of fluorescence intensity in all images was performed using Fiji software (https://imagej.net/Fiji).

12. ChIP-qPCR analysis

Two grams of 5-d-old etiolated Col-0 and *PIF3-myc* (in the Col-0 background) seedlings was cross-linked in 20 mL 1% (v/v) fresh formaldehyde solution under vacuum twice for 8 min each. To quench cross-linking, 2 M glycine was added to a final concentration of 0.125 M and was applied under vacuum for another 5 min. All vacuum steps were performed at –0.08 MPa. To protect RNA from degradation, 10 units per mL RNase inhibitor (Beyotime, R0102-KU) was added into buffers. Nuclei were pelleted by centrifugation at 4,000 g for 20 min at 4 °C, washed with nuclei extraction buffer 2 (10 mM Tris-HCl pH 8.0, 0.25 M sucrose, 10 mM $MgCl_2$, 1% [v/v] Triton X-100, 0.1 mM PMSF and protease inhibitor), and lysed in nuclei lysis buffer (50 mM Tris-HCl, pH 8.0, 10 mM EDTA, 1% [w/v] SDS, 0.1 mM PMSF and 1:100 (v/v) protease inhibitor cocktail [Bimake, Houston, USA]). Chromatin was sheared by sonication (220 W, 20 min, 4 s sonicating plus 9 s break, SCIENTZ IID) to approximately 500 bp. The chromatin solution was diluted 10-fold with ChIP dilution buffer (16.7 mM Tris-HCl, pH 8.0, 167 mM NaCl, 1.1% [v/v] Triton X-100, 1.2 mM EDTA, 0.1 mM PMSF and 1:100 (v/v) protease inhibitor cocktail [Bimake, Houston, USA]). Myc magnetic beads (Bimake, Houston, USA, 3 μL per IP) were washed with ChIP dilution buffer two times and then mixed with the chromatin solution and incubated at 4°C overnight. Immuno-complexes were precipitated and washed with four different buffers: low-salt buffer (20 mM of Tris-HCl, pH 8.0, 150 mM NaCl, 0.2% [w/v] SDS, 0.5% [v/v] Triton X-100, and 2 mM EDTA), high-salt buffer (20 mM Tris-HCl, 500 mM NaCl, 0.2% [w/v] SDS, 0.5% [v/v] Triton X-100, and 2 mM EDTA pH 8.0) , LiCl washing buffer（20 mM Tris-HCl, pH 8.0, 0.25 M LiCl, 1% [v/v] NP-40, 1% [w/v] sodium deoxycholate, and 1 mM EDTA pH 8.0) and TE washing buffer (10 mM Tris-HCl pH 8.0, and 1 mM EDTA pH 8.0). The bound chromatin fragments were eluted with the elution buffer (50 mM Tris-HCl, pH 8.0, 10 mM EDTA pH 8.0, and 1% [w/v] SDS) and the cross-linking was reversed by incubating at 65°C overnight. The mixture was treated with proteinase K for 1 h at 45°C to remove proteins. DNA was extracted with phenol/chloroform/isoamyl alcohol (25:24:1) and precipitated with two volumes of 100% ethanol at –80°C for 4 h. To recover the DNA, the sample was centrifugated at 16,000 g for 20 min at 4°C. The pellet was dried briefly and resuspended in 25 μL of TE buffer for further quantitative PCR analysis.

13. Electrophoretic mobility shift assay (EMSA)

Recombinant PIF3-GST protein and GST protein were used for EMSA. The probes used for the assays were synthesized and labeled with a fluorescein isothiocyanate (FITC) fluorescent probe (TsingKe Biological Technology) (Table S1). The DNA–PIF3 binding reaction consisted of 100 pg probe, 200 ng PIF3-GST protein, 10 mM Tris, pH 7.5, 5% (v/v) glycerol, 1 mM $MgCl_2$, 50 mM KCl, 0.2 mg/mL bovine serum albumin, 0.5 mM dithiothreitol, 0.5 mg/mL polyglutamate, and the indicated amount of unlabeled competitor. The reactions were incubated at room temperature for 30 min and fractioned by electrophoresis in a 6% native polyacrylamide gel (acrylamide:bisacrylamide, 29:1) containing 10% (v/v) glycerol, 89 mM Tris, pH 8.0, 89 mM boric acid, and 2 mM EDTA. The FITC signal was detected using a KODAK 4000MM Image Station.

14. Real-time PCR

RT-PCR was performed with gene-specific primers as listed in Table S1.

15. Measurement of the net $Ca^{2+}$ fluxes in the root cap

The NMT assay was performed as described previously[29]. Briefly, the net fluxes of $Ca^{2+}$ were measured non-invasively using the Non-invasive Micro-test Technology (NMT) with Non-invasive Micro-test system (NMS, NMT150S, Younger USA LLC, Amherst, MA, USA) and in Fluxes V2.0 (Younger USA LLC, Amherst, MA, USA) software. The microelectrode, LIX Holder, $Ca^{2+}$-LIX, and Ag/AgCl wire microsensor holder used in the experiment were purchased from Xuyue (Beijing) Science and Technology Co., Ltd., Beijing, China. In this study, plants were grown on medium for 3 days, before being transferred to the measurement solution (0.1 mM KCl, 0.1 mM $CaCl_2$, 0.1 mM $MgCl_2$, 0.5 mM NaCl, 0.3 mM MES, 0.2 mM $Na_2SO_4$, pH = 6.0). Calibration solution 1 (0.1 mM KCl, 0.02 mM $CaCl_2$, 0.1 mM $MgCl_2$, 0.5 mM NaCl, 0.3 mM MES, 0.2 mM $Na_2SO_4$, pH 6.0) and calibration solution 2 (0.1 mM KCl, 1 mM $CaCl_2$, 0.1 mM $MgCl_2$, 0.5 mM NaCl, 0.3 mM MES, 0.2 mM $Na_2SO_4$, pH 6.0) were used for correction. To avoid the interference caused by sloughing cells, the probe was positioned close to the lateral root cap instead of at the tip of the columella root cap, as shown in Fig. 4H. Ion flux recordings were taken for at least 8 min for each root cap and the measured data were rejected during the first minute. At least 4 plants were measured.

16. Yeast two hybrid (Y2H) assay

Y2H assays were performed as previously described[26,53]. Briefly, the coding sequences of *PIF3* were cloned in vector pGADT7 (AD). Primers for cloning are shown in Table S1. The sequence encoding the kinase domain of FER was cloned into pGBKT7 (BD). Distinct plasmid pairs were transformed into yeast AH109 cells. The transformants were diluted and plated onto synthetic defined (SD) medium lacking tryptophan and leucine (Trp–Leu) and SD medium lacking tryptophan, leucine, and histidine (–Trp–Leu–His) but containing with 20 mM 3-amino-1,2,4-triazole for 7 d to test the interaction.

17. Recombinant protein production in *Escherichia coli*

*PIF3* variants ($PIF3^{mut10A}$ and $PIF3^{mut10D}$) were synthesized by Qsingke Company. The coding sequence of *PIF3* or variants was cloned into the pGEX-4T-1 plasmid using restriction sites EcoR-I and BamH-I. The resulting constructs were transformed into *Escherichia coli* BL21 star (DE3) cells. Single clones were picked and transferred to LB medium for overnight culture at 37°C with shaking at 250 rpm. The cultures were diluted 1:50 with fresh LB medium containing 50 mg $mL^{-1}$ ampicillin for growth until reaching an optical density at 600 nm ($OD_{600}$) of 0.8. Protein production was induced with 0.5 mM isopropyl-β-D-thiogalactopyranoside (IPTG) at 16°C overnight. The cell cultures were spun down at 5,000 rpm for 8 min, and the cell pellet was resuspended in lysis buffer (25 mM Tris-HCl, pH 7.5, 300 mM NaCl, 20 mM imidazole, and 1 mM phenylmethylsulfonyl fluoride [PMSF]) and lysed with an ultrasonic cell disruptor (220 W, 20 min, 4 s sonicating plus 9 s break, SCIENTZ IID) on ice. The lysates were centrifuged at 12,000 g for 10 min at 4°C, and the resulting supernatant was transferred to a new tube and incubated with Ni-NTA resin at 4°C for 3 h. The Ni-NTA beads were washed extensively with wash buffer (20 mM Tris-HCl, pH 7.5, 300 mM NaCl, and 30 mM imidazole), and recombinant proteins were eluted with elution buffer (20 mM Tris-HCl, pH 7.5, 300 mM NaCl, and 300 mM imidazole). The eluates were desalted via centrifugation with a salt-free buffer (50 mM HEPES-NaOH, pH 7.5) using Amicon Ultra centrifugal filters (30K MWCO, Millipore). The centrifugation step was repeated five times to remove 99.9% of salts. The protein concentration was measured using a BCA kit (PC0020-500, Solarbio) according to the user manual. Aliquots of GST-PIF3, GST-PIF3$^{mut10A}$, and GST-PIF3$^{mut10D}$ proteins were then stored in 50 mM HEPES-NaOH, pH 7.5, at –80 °C until use.

18．Transgenic materials

For *Arabidopsis* transgenic lines, the coding sequences of *PIF3* variants (*PIF3*$^{mut10A}$ and *PIF3*$^{mut10D}$) variants were synthesized by Qsingke Company. *PIF3*, *PIF3*$^{mut10A}$ and *PIF3*$^{mut10D}$ were cloned were cloned into the *EcoR*I and *BamH*I restriction site of pCambia-130, upstreaming of the *Venus* sequence. These PIF3 variants were placed under the control of the *PIF3* promoter. *FER-GFP* was cloned into plant expression vector pDT7 with SpeI and PacI enzymes to generate the overexpression plants. Positive transformants were selected by resistance to basta or kanamycin and verified by western analysis using anti-GFP (Abclonal, AE012) or anti-Myc (Cell Signaling Technology, 2040). *FER-GFP* was crossed with *PIF3-myc* seedlings. Transgenic plants were generated by Agrobacterium (*Agrobacterium tumefaciens*)-mediated (GV3101 strain) transformation of *pif3-1*. Primary transformants ($T_1$) were selected based on hygromycin resistance, and $T_2$ lines containing only one T-DNA insertion were selected for further characterization by determining the Mendelian segregation ratio (3:1) of hygromycin-resistant seedlings in the $T_2$ progeny and confirmed in the $T_3$ progeny.

19. Co-immunoprecipitation assay

Seven-day-old *pFER::FER-GFP* transgenic seedlings were ground in liquid nitrogen, and the resulting tissue powder was resuspended in Co-IP buffer (50 mM Tris-HCl, pH 7.5, 150 mM NaCl, 1 mM EDTA, 5% [v/v] glycerol, and protease inhibitor mixture containing 1% [v/v] Triton X-100, 1 mM PMSF and 50 μM MG132). The homogenate was centrifuged at 12,000 g for 10 min. Then supernatant was incubated with glutathione-agarose preincubated with PIF3-GST or GST for testing the interaction between FER and PIF3. The incubation was carried out for 2 h at 4°C, and then the agarose beads were washed three times with the Co-IP buffer containing 0.1% Triton X-100. The immunoprecipitates associated with the agarose beads were boiled in SDS/PAGE loading buffer, analyzed by SDS/PAGE, and transferred to nitrocellulose membrane for detection with anti-GFP (AE012, ABclonal Technology) and anti-GST (AE001, ABclonal Technology) antibodies. For the *in vivo* Co-IP, 5-d-old etiolated *PIF3-myc FER -GFP* seedlings were used to lysate as above. Myc magnetic beads (Bimake, Houston, USA, 50 μL per IP) were washed with Co-IP buffer two times and then mixed with the lysate and incubated at 4°C overnight. Anti-GFP (AE012, ABclonal Technology) and anti-myc (Cell Signaling Technology, 2040) were used for the immunoblot assay.

20. GST pull-down assay

The recombinant His-FER-KD and GRP7-GST proteins were co-incubated with 50 μL of glutathione-agarose beads (Thermo Fisher Scientific, 16100) in binding buffer (50 mM Tris-HCl pH 8.0, 150 mM NaCl, and 10 mM $MgCl_2$) at 4°C for 6 h. The beads were washed with washing buffer I (50 mM Tris-HCl, pH 8.0, 300 mM NaCl, and 0.5% (v/v) Triton X-100) for 10 min and then two times with washing buffer II (50 mM Tris-HCl, pH 8.0, 150 mM NaCl, and 10 mM $MgCl_2$), each for 20 min. The proteins on the beads were eluted by boiling in SDS loading buffer, separated by SDS-PAGE, and detected by immunoblotting using an anti-GST (SC-80998, Santa Cruz Biotechnology) or anti-His antibody (M20001, Abmart).

21. *In vitro* phosphorylation assay

An *in vitro* phosphorylation assay was performed as previously described[53], but with minor modifications. Recombinant FER-KD and FER-KD$^{K565R}$ mutant proteins that co-expressed with λ-PP were purified. The FER-KD or FER-KD mutant proteins (1 μM) were co-incubated with recombinant PIF3-GST at room temperature for 40 min in 50 μL of assay kinase buffer containing 1 mM ATP and 10 mM $Mg^{2+}$. Proteins were resolved by SDS-PAGE and transferred to a nitrocellulose membrane (PALL, P-N66485) for protein immunoblotting using anti-phosphothreonine (Abcam, ab218195), anti-phosphoserine (Abcam, ab9332), and anti-phosphotyrosine (Abcam, ab179530) antibodies. The proteins were visualized by immunoblotting.

22. Identification of phosphorylation sites

After phosphorylation by FER-KD, the PIF3-GST bands were excised and cut into small pieces after Coomassie Brilliant Blue staining. Then, the pieces were dehydrated, and the proteins were reduced, alkylated, digested, and analyzed by mass spectrometry (MS) as described previously[56]. For label-free quantification, raw files were processed using MaxQuant (version 1.6.1.0) and Proteome Discoverer (Thermo Fisher Scientific, version 1.4). PIF3 phosphopeptides were identified by searching all tandem MS spectra against the Araport11_ pep_201606 sequence database and filtering using a FDR of < 0.01 at the peptide level and < 0.05 at the protein level by MaxQuant. We selected carbamidomethylation of cysteines as the fixed modification and used oxidation (M) and phosphorylation (STY) as variable modifications. In addition, we used Proteome Discoverer to assess phosphorylation localization. After identification, we performed PIF3 protein label-free quantitation of phospho-peptides with the MaxLFQ algorithm integrated in the MaxQuant software suite.

23. *In vivo* PIF3 abundance analysis

Seeds of Col-0 and *PIF3-myc* were surface sterilized, stratified, and sown on agar plates containing solidified half-strength MS medium before being placed vertically at 22°C in long-day conditions for 2 d. The seedlings were moved in the dark for 3 d, and incubated with half-strength liquid MS containing 1 μM synthesized RALF1 for 5h. Approximately 1 g of seedlings was ground in liquid nitrogen and then incubated with Co-IP buffer (50 mM Tris-HCl, pH 7.5, 150 mM NaCl, 1% [v/v] NP-40, 0.25% [w/v] sodium deoxycholate, 5% [v/v] PPVP, 10% [v/v] glycerol, and 10 mM $MgCl_2$). The PIF3-myc protein was visualized by immunoblotting.

24. *In vitro* degradation assay for PIF3-related recombinant proteins

Total proteins were extracted by homogenizing the roots of 7-day-old Col-0 plants in the presence of 150 mM NaCl, 12.5 mM citric acid, 50 mM $Na_2HPO_4$, and pH 6.5 (500 μL of extraction buffer per g tissue). The homogenate was incubated for 1.0 h on ice. The homogenate was centrifuged at 12,400g for 10 min at 4°C, and the supernatant was collected. This centrifugation step was repeated twice. Protein concentration was determined in supernatants using the Bradford protein assay method (Bradford reagent; Sigma-Aldrich). Equal amounts of protein samples (2 mg total proteins) were mixed with 200 μg GST-PIF3, GST-PIF3$^{mut10A}$ and GST-PIF3$^{mut10D}$ proteins. Mixtures were incubated at 22°C for 5 min. The GST-tagged proteins were visualized by immunoblot. The gray value of the bands was used to calculate the relative abundance of protein. The quantification of gray value was performed using Fiji software (https://imagej.net/Fiji).

### Acknowledgements

The authors are grateful to Meixia Hu and Shufeng Song (State Key Laboratory of Hybrid Rice, Hunan Hybrid Rice Research Center) for their technical support on confocal microscope. We thank Dr. Hongdong Chen (College of Life Sciences, Peking University) for the kind gift of *PIF3-GFP* seeds. Weijun Chen and other members in the laboratory are thanked for critical reading the manuscript and helpful discussions. The sequencing was conducted by OE Biotech Co., Ltd. (Shanghai). The work was supported by grants from National Natural Science Foundation of China (NSFC-32070769), and China Postdoctoral Science Foundation funded project (2020M672475), and the Science and Technology Innovation Program of Hunan Province (2021JJ40060). **Authors' contributions**: F. Y. and F. X. designed the research; F. X., J. C., Y.B. L., S.L.O., Y.R.W., and X.M.F. performed the experiments; F. X. F. Y. and K.H. wrote the manuscript. All authors reviewed and approved the manuscript for publication.

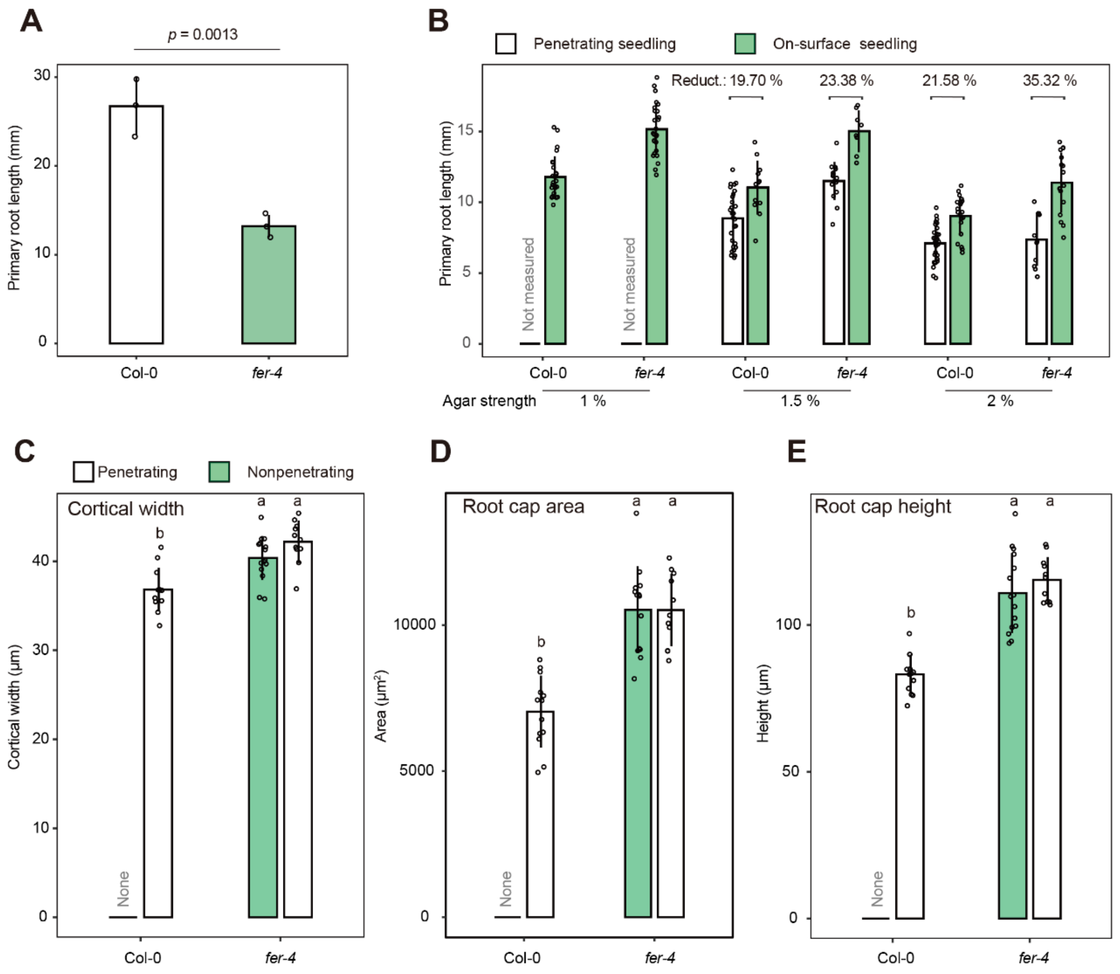


**Extended Data Fig. 1 Primary root length of *fer-4* growing on sand or solidified MS medium.**

A. Primary root length of 10-d-old Col-0 and *fer-4* seedlings growing on sand (*P*-value was obtained by Student's t-test comparing to Col-0 to *fer-4*). B. Primary root length of Col-0 and *fer-4* seedlings penetrating agar-solidified MS medium (penetrating seedling) or growing on the surface of MS medium (On-surface seedling). Due to the appearance of helical roots on 1% agar-solidified medium, the primary root length could not be measured. C-E. Cortical width (C), root cap area (D), and root cap height (E) of Col-0 and *fer-4* roots that could (penetrating) or could not (nonpenetrating) penetrate into agar-solidified MS medium. Data are means ± SD; for (C-E), data were analyzed by one-way ANOVA with Tukey's test. Different lowercase letters indicate significant differences ($P < 0.05$).

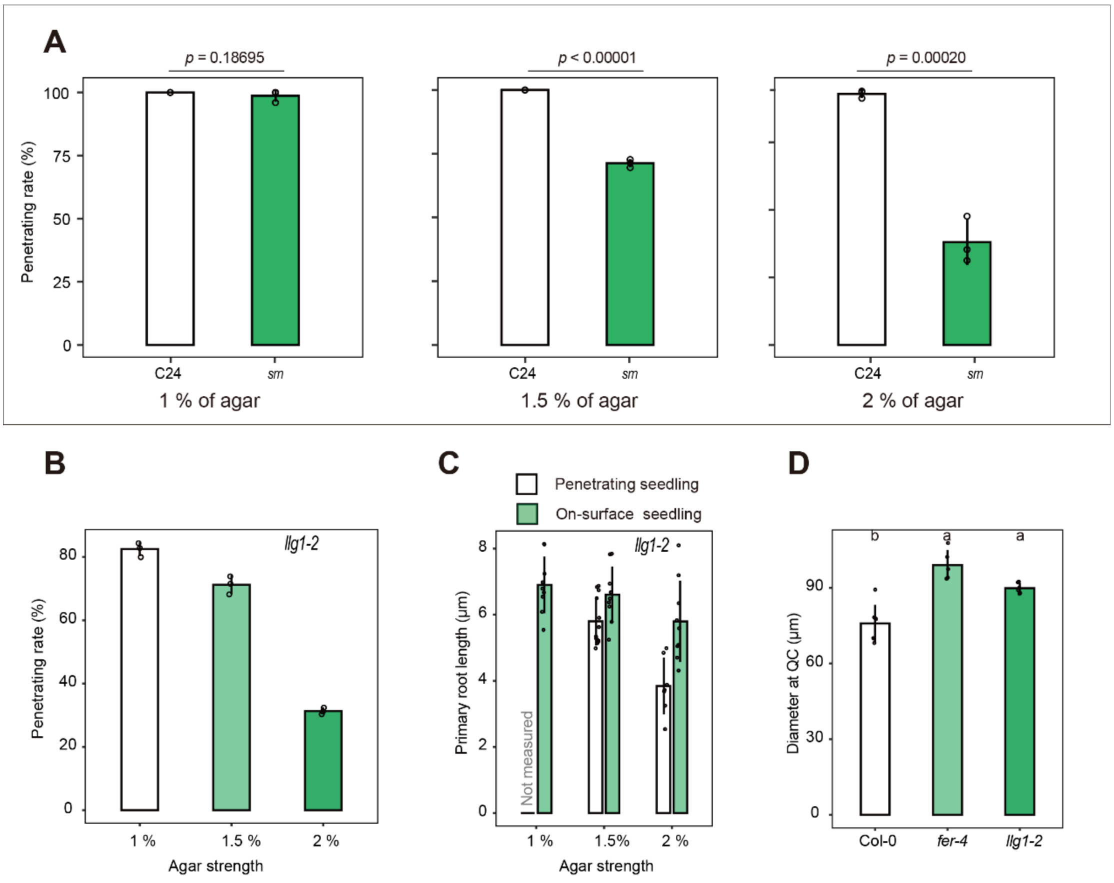


**Extended Data Fig. 2 Phenotyping of *srn* and *llg1-2* mutants in the penetration assay.**

A. Rate of root penetration of the wild-type C24 and the *srn* mutant in medium solidified with different agar concentrations. Data are means ± SE (n = 3 plates, more than 20 seedlings per agar concentration). The *P*-values were obtained by Student's *t*-test comparing to C24 to *srn.* B. Rate of root penetration of the *llg1-2* mutant in the medium solidified with different agar concentrations. C. Primary root length of *llg1-2* seedlings penetrating in agar-solidified MS medium (penetrating seedling) or growing on the surface of the MS medium (On-surface seedling). Due to the appearance of helical roots on medium solidified with 1% agar, the primary root length could not be measured. D. Diameter of *fer-4* and *llg1-2* roots at the quiescent center (QC). Data are mean ± SD. Data were analyzed by one-way ANOVA with Tukey's test. Different lowercase letters indicate significant differences ($P < 0.05$).

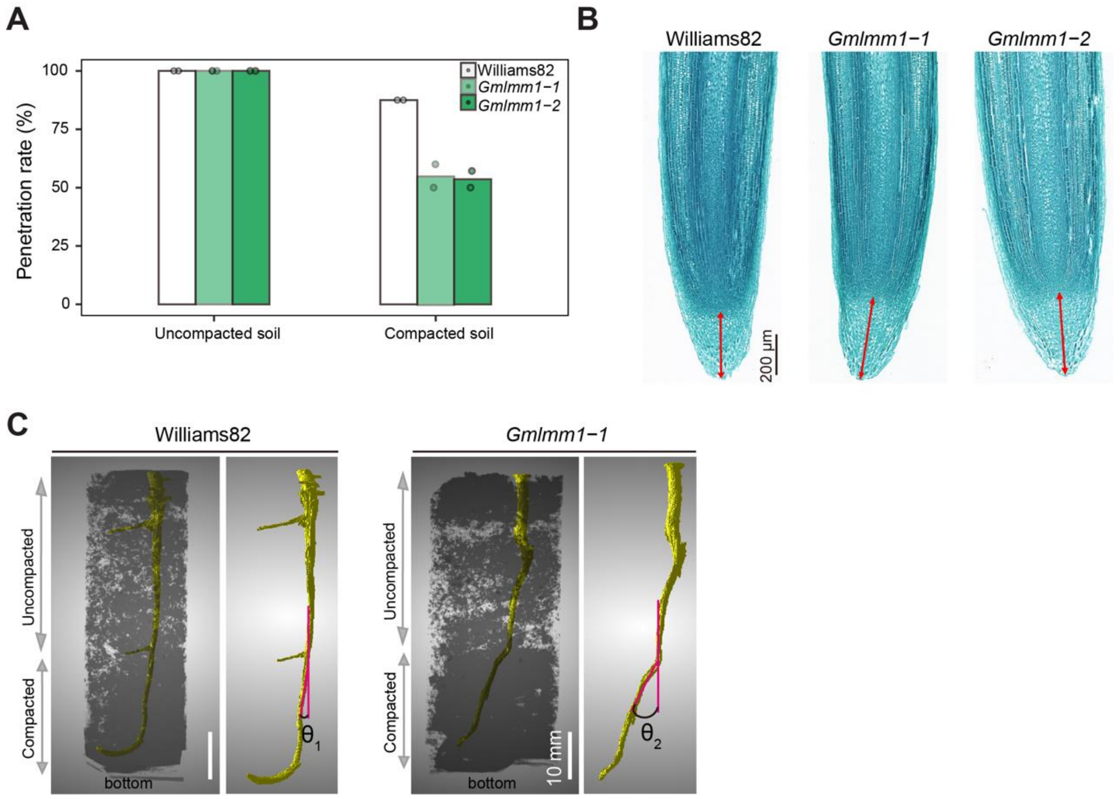


**Extended Data Fig. 3. Soybean *fer*-related mutants in penetration assay and their root cap phenotype.**

A. Soybean root penetration rate in uncompacted (1.0 g cm$^{-3}$) and compacted (1.8 g cm$^{-3}$) soil. n>7 from two biological replicates. B. *Gmlmm1-1* and *Gmlmm1-2* have a taller root cap than Williams 82. The roots of 5-d-old soybean seedlings grown in soil were used for anatomical slice. The red lines with double arrows indicate the height of the root cap. C. CT images showing soybean root growing in compacted soil. θ indicates the bending angle of root when penetrating into the compacted soil (1.4 g cm$^{-3}$).

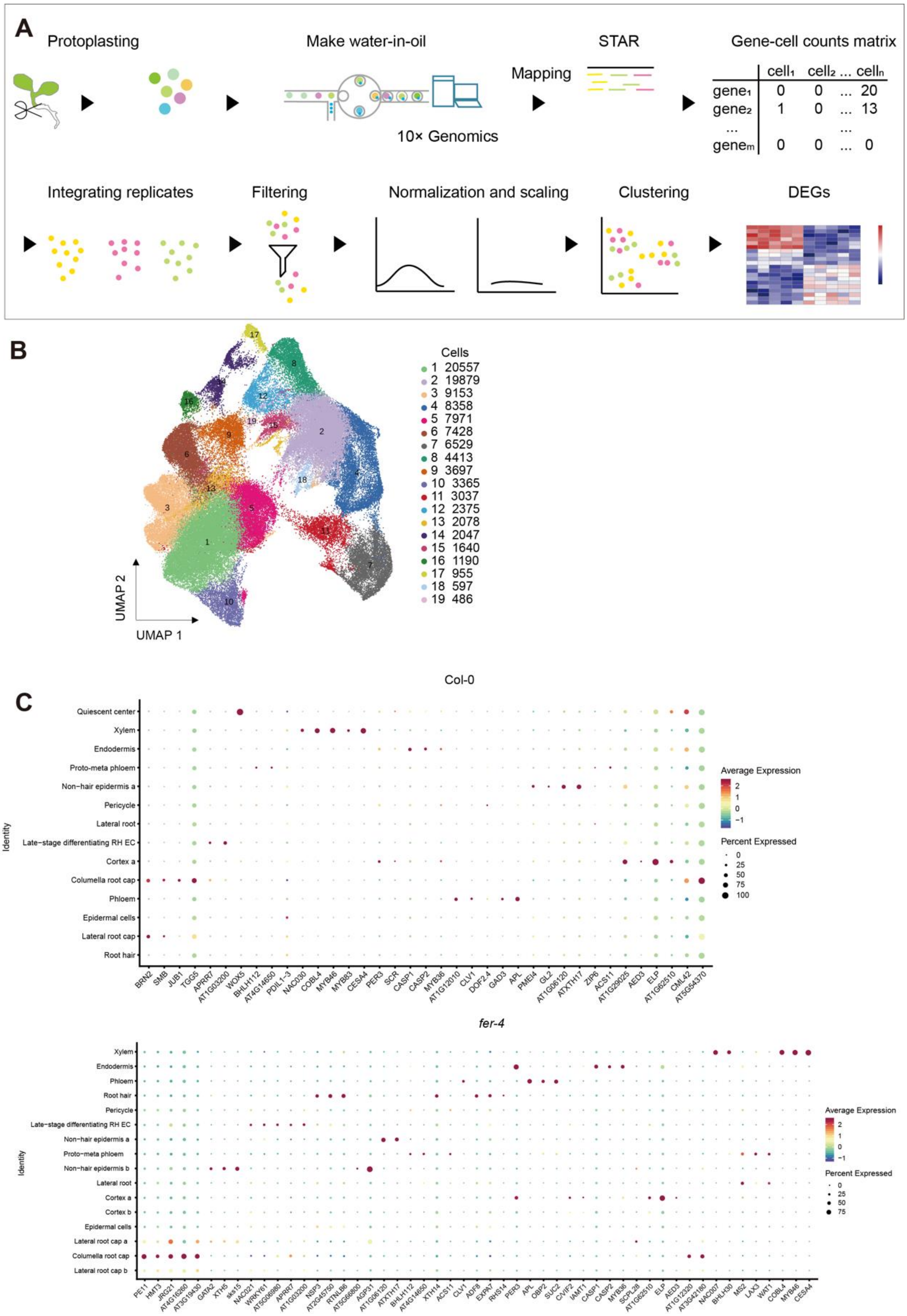


**Extended Data Fig. 4 Pipeline and tissue-specific reporter genes used in this study.**

A. Overview of the scRNA-seq pipeline, including metrics of the scRNA-seq experiment. B. UMAP visualization to identify 19 putative cell clusters from 12 root samples of Col-0 and *fer-4*. Each dot denotes a single cell. Colors denote corresponding cell clusters. C. Dot-plot of identified tissue specific genes for each cell identity cluster, containing known and uncharacterized genes. The size of the circles represents the percentage of cells with expression (pct.exp.), while the color indicates the scaled average expression level of this gene in each cell type (avg exp. scale).

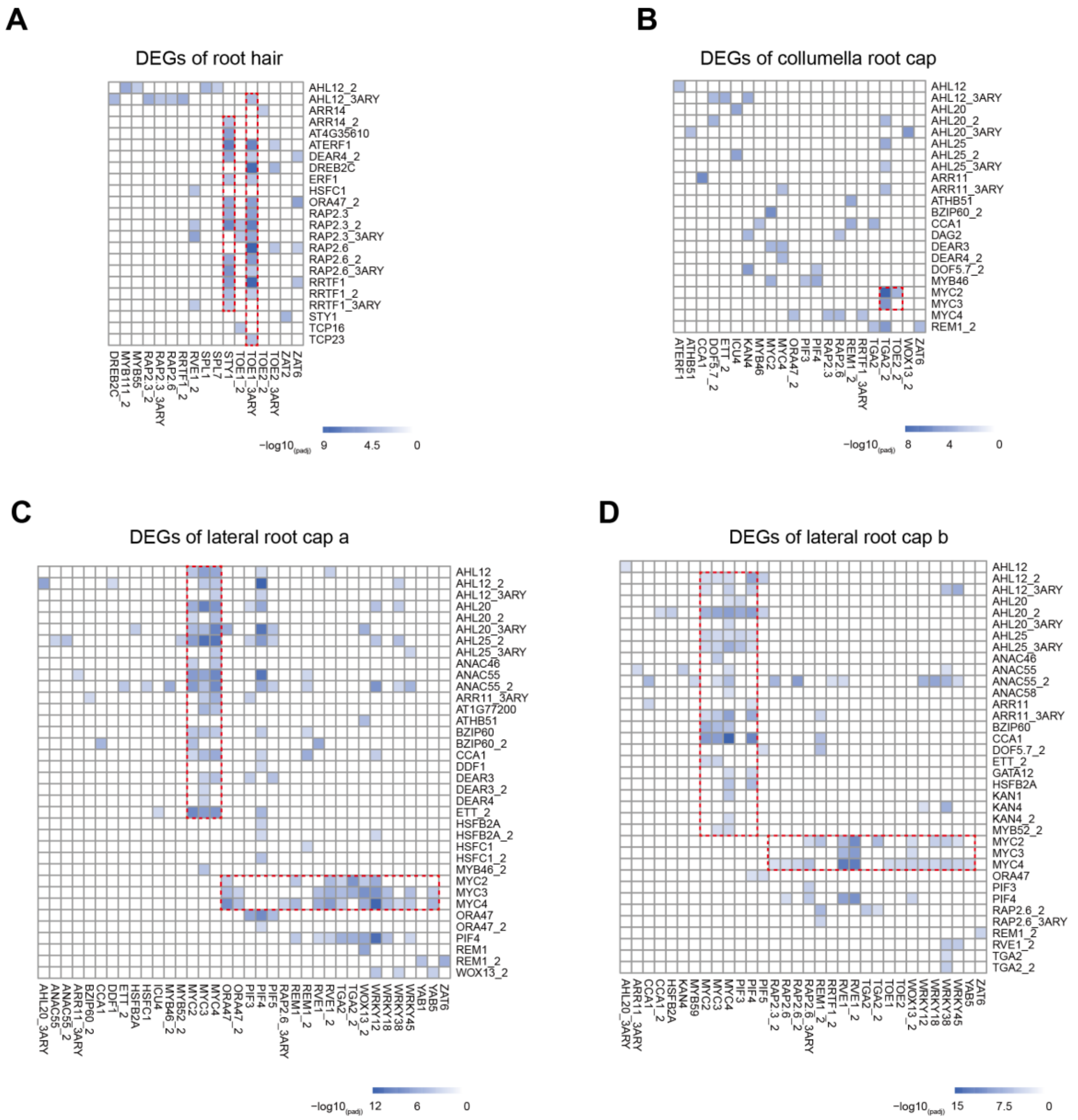


**Extended Data Fig. 5 Analysis of transcription factor genes that are regulated by FER using PMET.**

A-D. Heatmaps representing *P*-values ($Log10_{Padj}$) for enrichment of motif pairs in root hair (A), columella root cap (B), and lateral root cap a (C) and b (D) identity genes. The x- and y-axes of all heatmaps show the same order of transcription factor binding motifs. Pixel colors indicate significance values ($Log10_{Padj}$) for the abundance of the given motif pairs in the promoters of identity genes per cell type.

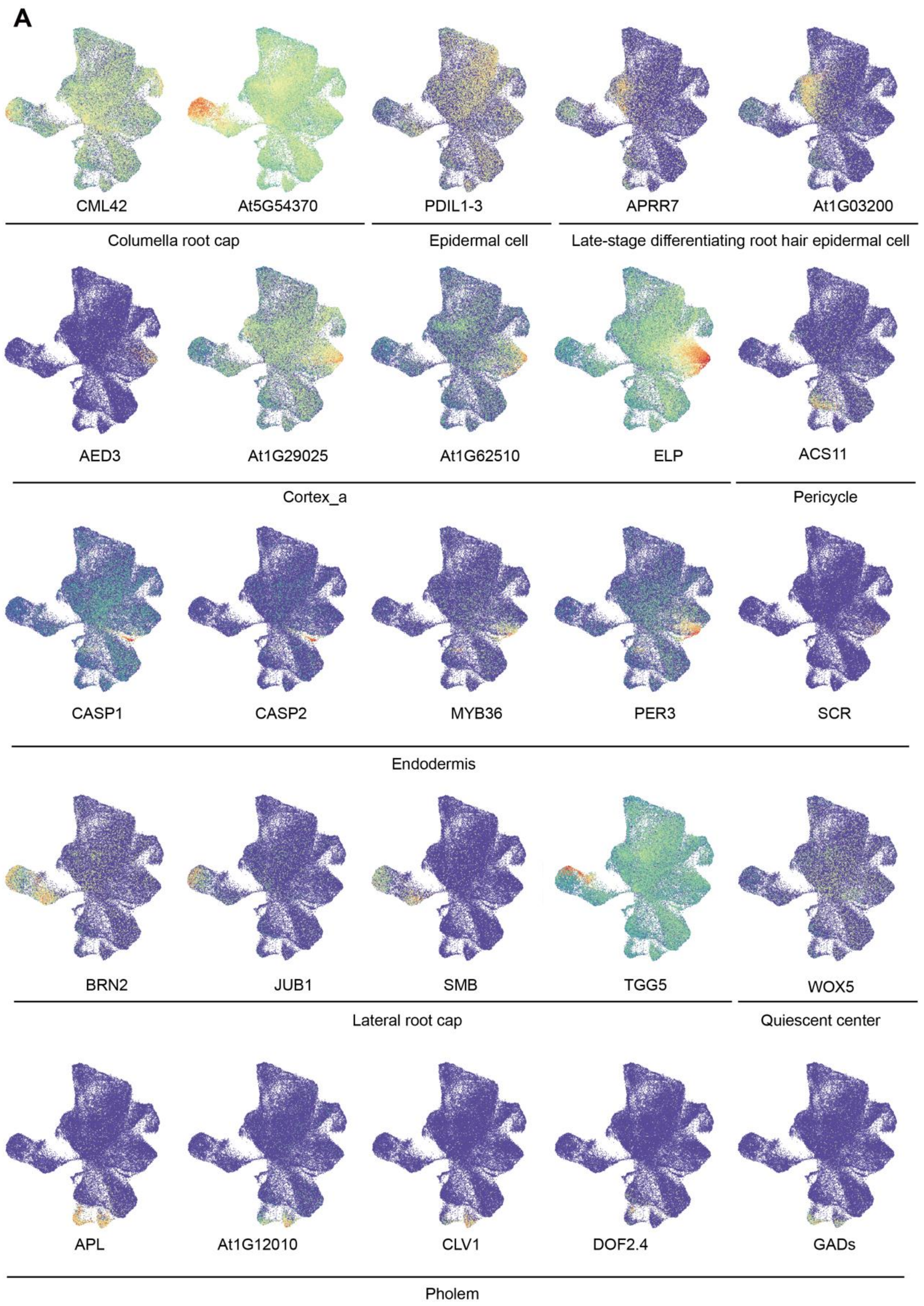
A
CML42
At5G54370
PDIL1-3
APRR7
At1G03200
Columella root cap
Epidermal cell
Late-stage differentiating root hair epidermal cell
AED3
At1G29025
At1G62510
ELP
ACS11
Cortex_a
Pericycle
CASP1
CASP2
MYB36
PER3
SCR
Endodermis
BRN2
JUB1
SMB
TGG5
WOX5
Lateral root cap
Quiescent center
APL
At1G12010
CLV1
DOF2.4
GADs
Pholem

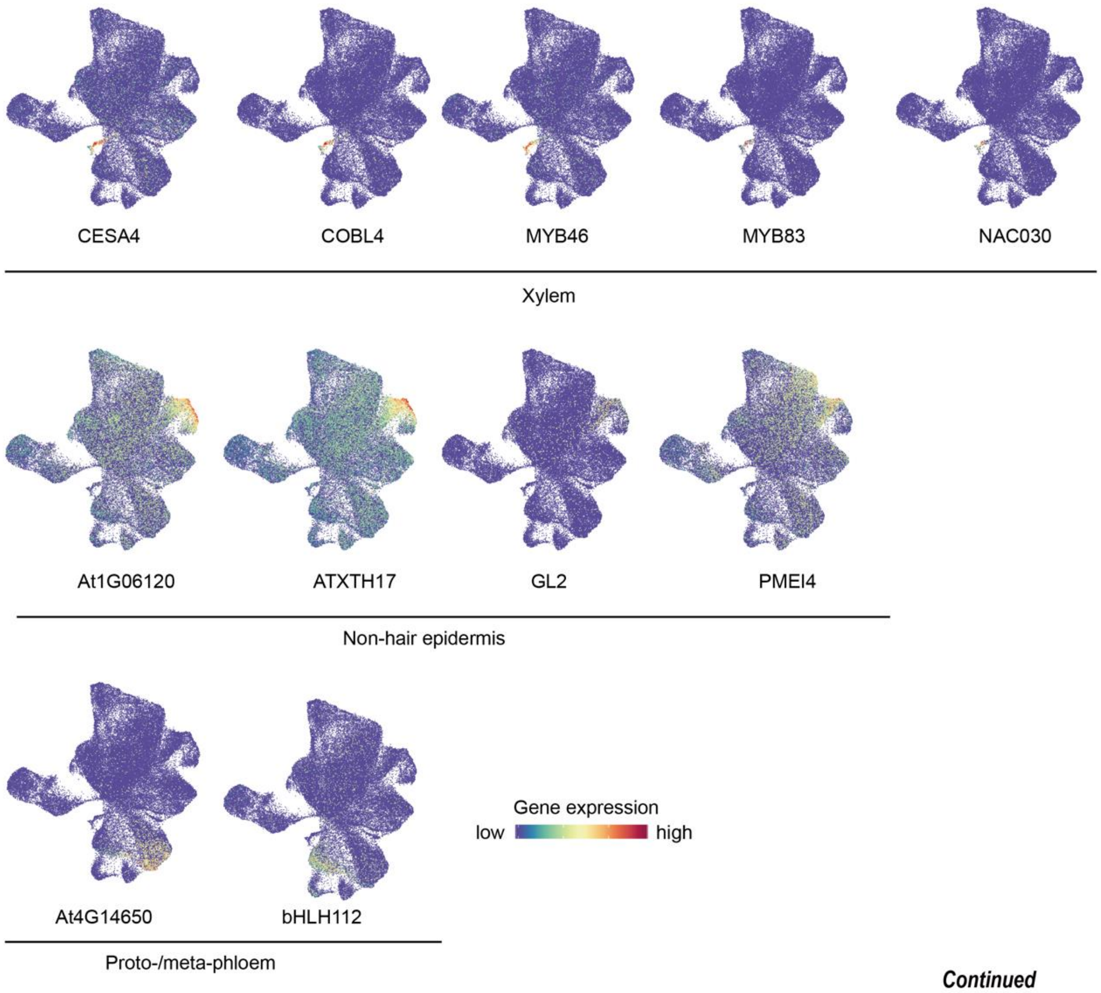
CESA4
COBL4
MYB46
MYB83
NAC030
Xylem
At1G06120
ATXTH17
GL2
PMEI4
Non-hair epidermis
At4G14650
bHLH112
Proto-/meta-phloem
Gene expression
low
high

***Continued***

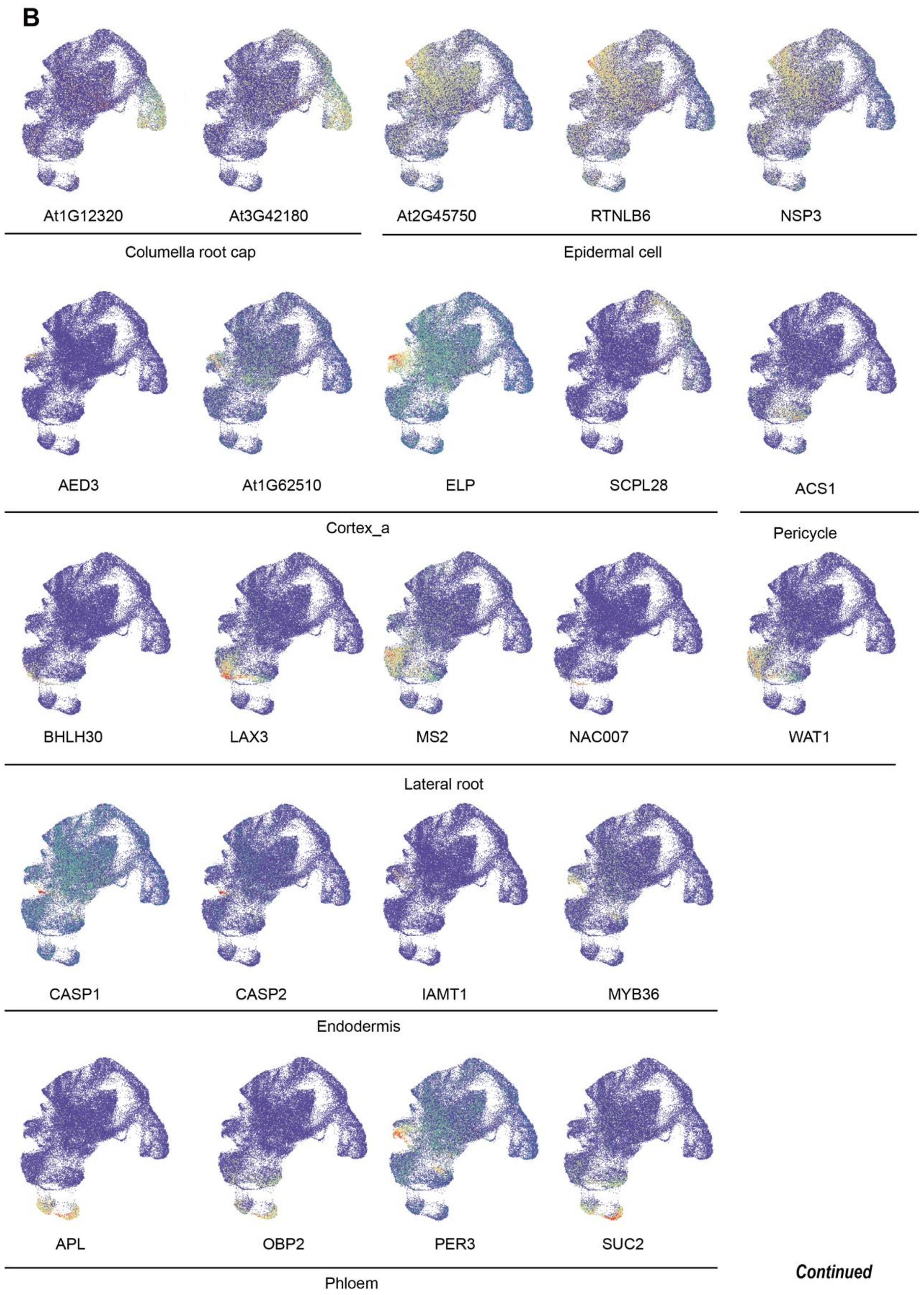
B
At1G12320
At3G42180
At2G45750
RTNLB6
NSP3
Columella root cap
Epidermal cell
AED3
At1G62510
ELP
SCPL28
ACS1
Cortex_a
Pericycle
BHLH30
LAX3
MS2
NAC007
WAT1
Lateral root
CASP1
CASP2
IAMT1
MYB36
Endodermis
APL
OBP2
PER3
SUC2
Phloem

Continued

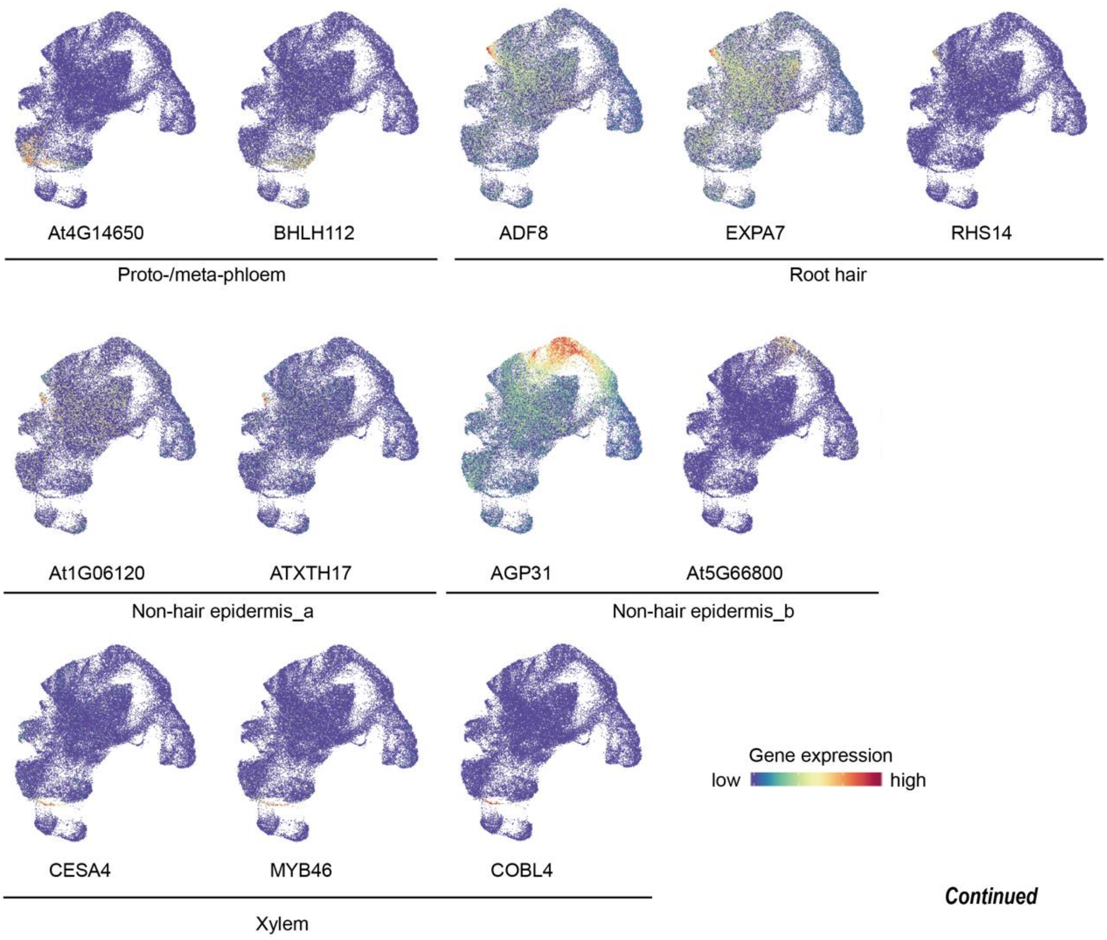


*Continued*

**Extended Data Fig. 6 Expression of marker genes.**

A and B. UMAP plots of Col-0 (A) and *fer-4* (B) showing the selected marker genes for different cell types. The full names and referenced expression patterns of selected genes are summarized.

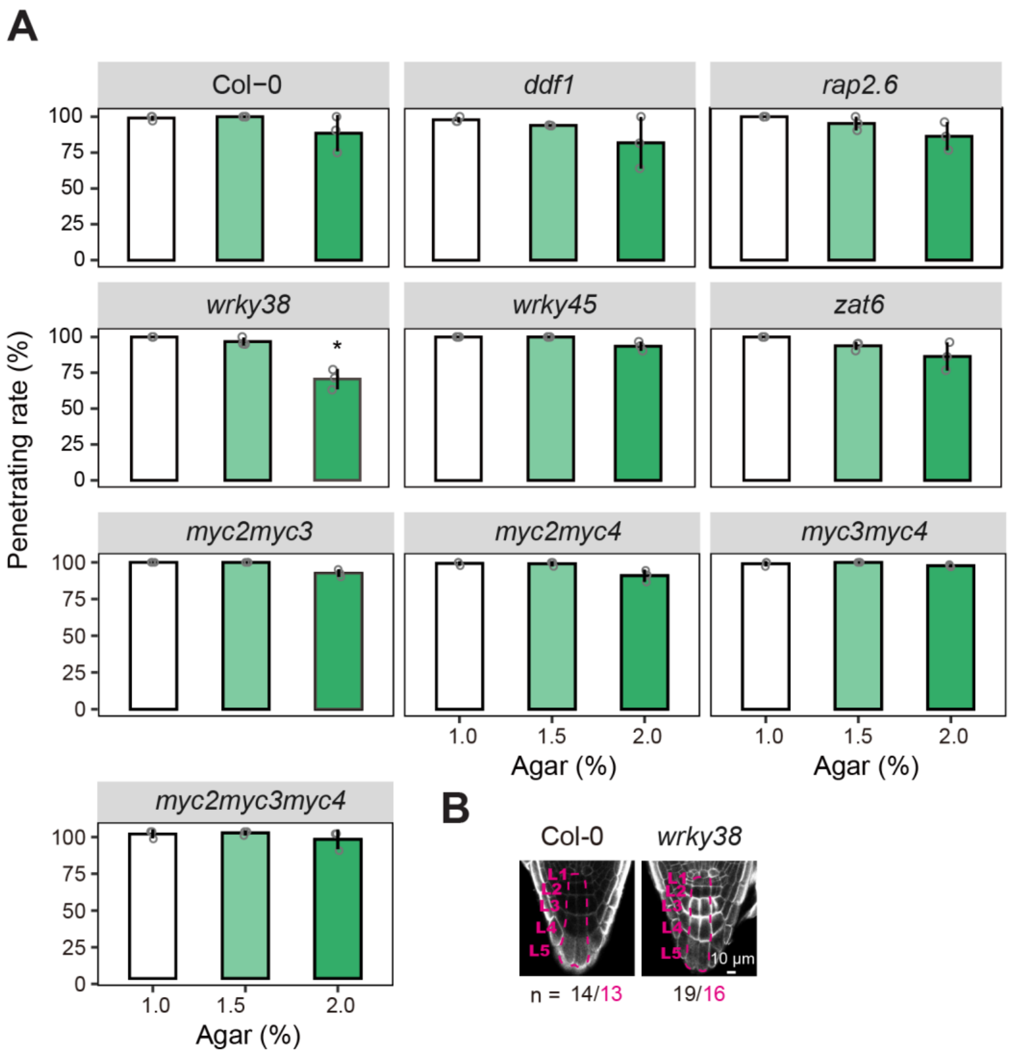


**Extended Data Fig. 7 Root penetration rate of the mutants for selected transcription factors that were enriched using PMET and iGRN methods.**

A. Root penetration rate of Col-0 and the *ddf1*, *rap2.6*, *wrky38, wrky45*, *zat6*, *myc23*, *myc24*, *myc34* and *myc234* mutants in medium solidified with different agar concentrations. These transcription factors are also listed in Figure 3D. Data are means ± SE (n = 3 plates, more than 20 seedlings per agar concentration). Significant differences were determined by one-way ANOVA with Tukey's test (*$P < 0.05$) between the mutant and Col-0 at same agar concentration. B. Root cap phenotype of *wrky38*. The dashed magenta line indicates the columella root cap (CRC) cell. L: layer.

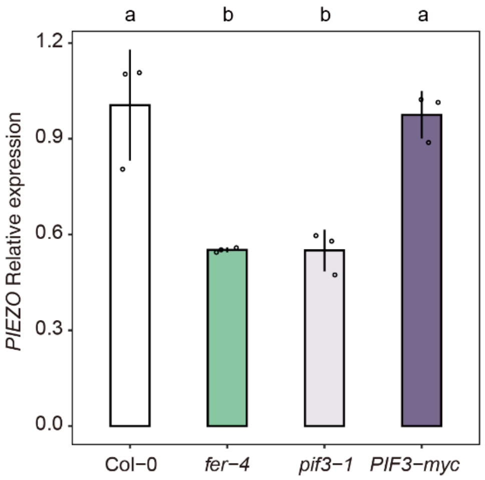


**Extended Data Fig. 8 *PIEZO* expression in roots.**

RT-PCR analysis of *PIEZO* transcript in the root of Col-0, *fer-4, pif3-1*, and *PIF3-myc*. *ACTIN* was used as an internal control for quantification. Data are means ± SE (n = 3).

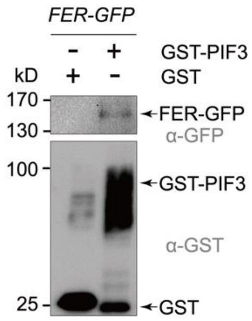


**Extended Data Fig. 9 Co-IP assay performed with crude extracts of *FER-GFP* transgenic plants.**

FER-GFP associates with recombinant GST-PIF3, but not with GST, when subjected to a CO-IP assay. Immunoprecipitated GST-PIF3 and coimmunoprecipitated FER-GFP were detected using anti-GST and anti-GFP, respectively. GST was used as negative controls.

416

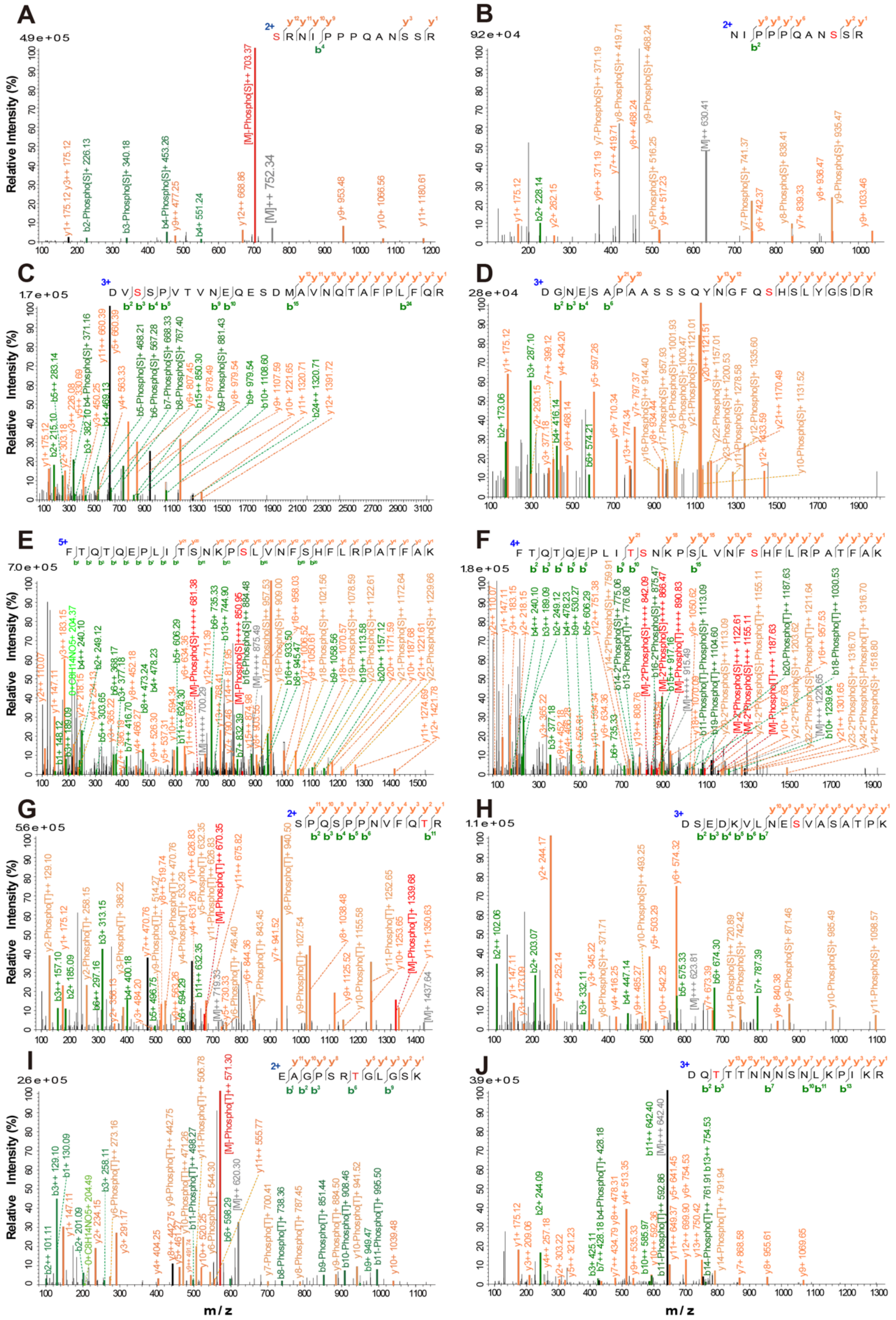

417 **Extended Data Fig. 10 ESI mass spectrometric analysis of PIF3 phosphorylated residues.**

Identified PIF3 phosphorylated residues were Ser-48 (A), Ser-58 (B), Ser-115 (C), Ser-160 (D), Ser-196 (E), Ser-201 (F), Thr-234 (G), Ser-250 (H), Thr-335 (I), and Thr-499 (J). The identified peptide sequences are displayed. The y-ion and b-ion are shown above the sequences.

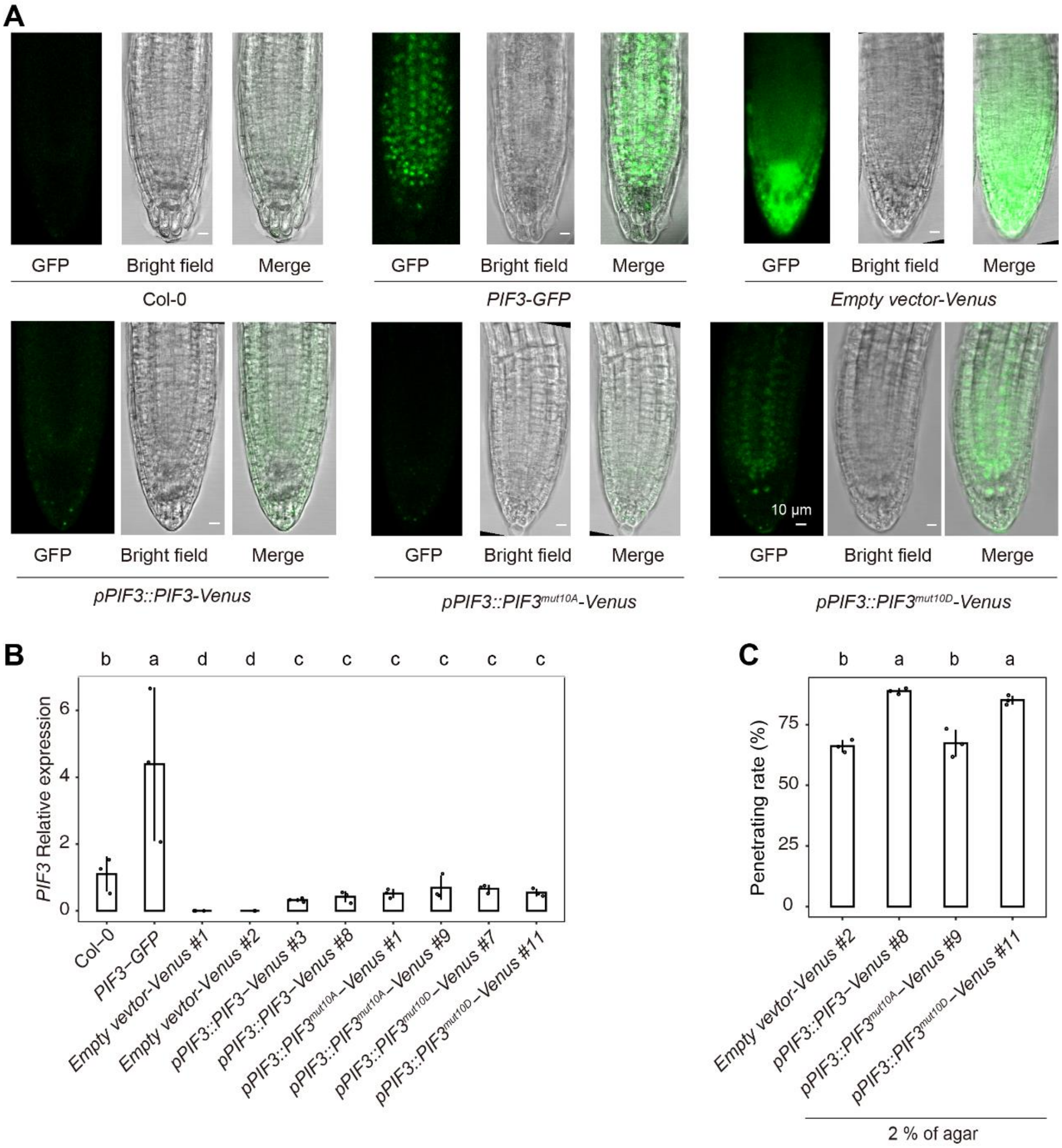


**Extended Data Fig. 11 PIF3 proteins accumulated in the nucleus of the root cap cell.**

A. GFP fluorescence intensity and expression pattern of Col-0, *35S::PIF3-GFP in Col-0* (*PIF3-GFP*), P1300-LV empty vector in *pif3-1* (*Empty vector-Venus*), *pPIF3::PIF3-Venus*, *pPIF3::PIF3*$^{mut10A}$*-Venus*, and *pPIF3::PIF3*$^{mut10D}$*-Venus* five-day-old etiolated seedlings grown in dark. Root tips were observed using same setups of confocal microscope. B. RT-PCR analysis of *PIF3* transcript in seedlings of Col-0, *PIF3-GFP, Empty vector-Venus*, *pPIF3::PIF3-Venus*, *pPIF3::PIF3*$^{mut10A}$*-Venus*, and *pPIF3::PIF3*$^{mut10D}$*-Venus*. ACTIN was used as an internal control for quantification. Data are means ± SE (n = 3). C. Behavior of *Empty vector-Venus*, *pPIF3::PIF3-Venus*, *pPIF3::PIF3*$^{mut10A}$*-Venus*, and *pPIF3::PIF3*$^{mut10D}$*-Venus* seedlings in the penetration assay. For B and C, different lowercase letters indicate significant differences at $P < 0.05$.

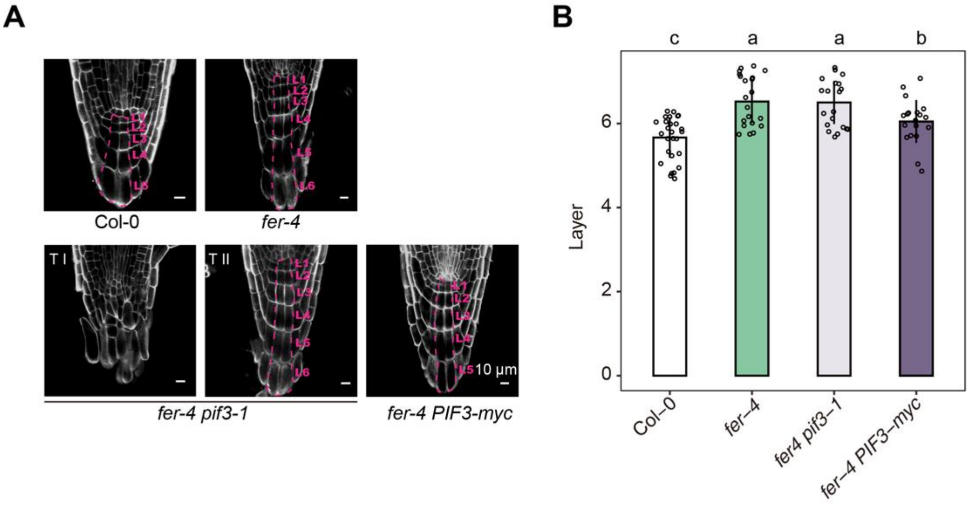


**Extended Data Fig. 12 Overexpression of PIF3 partially rescues the defect in root cap of *fer-4*.**

A. Root cap phenotype of Col-0, *fer-4*, *fer-4 pif3-1*, and *fer4 PIF3-myc*. The dashed magenta line indicates the columella root cap (CRC) cells. L: layer. For *fer-4 pif3-1* double mutant, two types (~19.0% T I and 81.0% T II) of root cap were found. B. Cell layer number of in the CRC of Col-0, *fer-4*, *fer-4 pif3-1*, and *fer4 PIF3-myc*. Data were analyzed by one-way ANOVA with Tukey's test. Different lowercase letters indicate significant difference ($P < 0.05$).

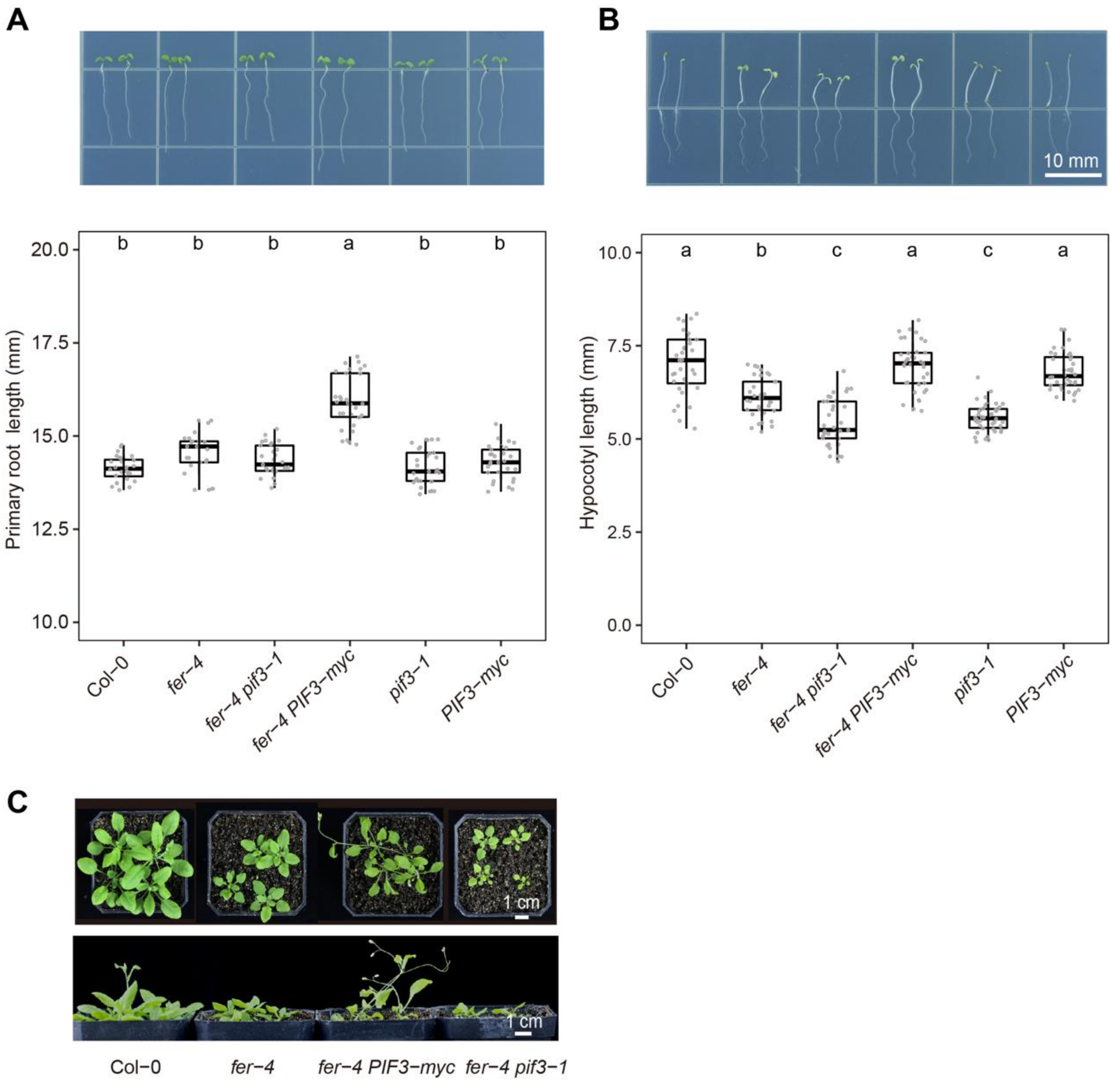


**Extended Data Fig. 13 Overexpression of PIF3 altered the phenotypes of *fer-4*.**

A. Primary root length of 5-d-old seedlings of Col-0, *fer-4*, *fer-4 pif3-1*, *fer4 PIF3-myc, pif3-1* and *PIF3-myc*. B. Hypocotyl length of 5-d-old etiolated seedlings of each genotype. C. Representative images of each genotype grown in soils for 5 weeks under long-day conditions (16h light / 8h dark). For A and B, data were analyzed by one-way ANOVA with Tukey's test. Different lowercase letters indicate significant difference ($P < 0.05$).
